# 2D-HELS-AA MS Seq: Direct sequencing of tRNA reveals its different isoforms and multiple dynamic base modifications

**DOI:** 10.1101/767129

**Authors:** Ning Zhang, Shundi Shi, Xuanting Wang, Wenhao Ni, Xiaohong Yuan, Jiachen Duan, Tony Z. Jia, Barney Yoo, Ashley Ziegler, James J. Russo, Wenjia Li, Shenglong Zhang

## Abstract

We report a direct method for sequencing tRNA^Phe^ without cDNA by combining 2-dimensional hydrophobic RNA end-labeling with an anchor-based algorithm in mass spectrometry-based sequencing (2D-HELS-AA MS Seq). The entire tRNA^Phe^ was sequenced and the identity, location and abundance of all 11 base modifications were determined. Changes in ratios of wybutosine and its depurinated form under different conditions were quantified, pointing to the ability of our technology to determine dynamic changes of nucleotide modifications. Two truncated isoforms at 3’CCA tail of the tRNA^Phe^ (75 nt CC, 80% and 74 nt C, 3%) were identified in addition to the 76 nt tRNA^Phe^ with a full-length 3’CCA tail (17%). We also discovered a new isoform with A-G transitions at both the 44 and 45 positions in the tRNA^Phe^ variable loop.

**One Sentence Summary:** Direct 2D-HELS-AA MS Seq of tRNA reveals different isoforms and base modifications

---

As an essential component of protein synthesis machinery, transfer RNA (tRNA) is present in all living cells and more than 600 different tRNA sequences and a large breadth of different post-transcriptional base modifications that have been reported (*1, 2*). Despite its significance, structural and functional studies to understand the underlying biochemistry of tRNA itself have been hindered due to the lack of efficient tRNA sequencing methods. Although the first tRNA was sequenced, thanks to a herculean effort, by Holley in 1965 (*3*), tRNAs are currently the only class of small cellular RNAs that cannot be efficiently sequenced with current sequencing techniques (*4*). Typically, methods used to sequence tRNAs are indirect and require complementary DNA (cDNA) intermediates. However, cDNA synthesis results in a loss of endogenous base modification information originally carried by RNAs, resulting in the inability to accurately sequence the rich and dynamic base modifications in tRNAs which are an intrinsic part of their structure and function. Other sequencing methods that do not involve cDNA can detect base modifications, but these techniques usually require harsh sample treatments such as intensive enzymatic or chemical hydrolysis, resulting loss of in spatial modification information (*5-7*). As opposed to indirect sequencing methods, direct sequencing of RNAs without cDNA synthesis would theoretically allow identification of all associated modified nucleotides in any RNA sample.

One potential direct RNA sequencing method is mass spectrometry (MS)-based sequencing, which directly correlates the mass of each base, rather than proxies such as fluorescence (*8*) or fluctuations in electric current (*9*) to sequence outputs. The four canonical ribonucleotides and most modified bases either have inherently unique masses or can be easily converted into different signature masses, allowing them to potentially be used for *de novo* MS-based direct RNA sequencing. Previously, top-down MS and tandem MS (*10–12*) have been used in attempts to directly sequence tRNAs. However, these traditional MS methods have significant methodological inadequacies in the preparation of high-quality sequence ladders and subsequently in the reduction of complicated data, resulting in inaccurate base-calling (*13, 14*). Therefore, a two-dimensional (2D) LC-MS-based RNA sequencing method was established to produce easily-identifiable mass-retention time (t_R_) ladders (*14*), allowing *de novo* sequencing of short single-stranded RNAs. We recently introduced a hydrophobic end-labeling strategy (HELS) to 2D LC-MS-based RNA sequencing by introducing 2D mass-t_R_ shifts for ladder identification of mixed RNAs (*15*), allowing both the complete reading of a sequence from a single ladder and the sequencing of mixed RNA samples. Despite its success in model studies on short RNAs (~ 20 nt) (*15*), it is still challenging to perform *de novo* MS sequencing of biological RNA such as tRNAs due to the data complexity caused by the increased length (typically ~76-90 nt) and abundant modifications. To simplify the 2D HELS data analysis, here we developed two computational anchor algorithms which innovatively accomplish automated sequencing of RNAs. The signature t_R_-mass value of the hydrophobic tag specifies the exact starting data point, the anchor, for the algorithm to accurately determine data points corresponding to the desired ladder fragments, significantly simplifying data reduction and enhancing the accuracy of sequence generation. The idea of using an anchor to identify sequence ladder start-points can be generalized and extended to any known chemical moiety beyond hydrophobic tags, *e.g.*, PO4^-^ at the beginning of the tRNA or any nucleotide with a known mass. We can program its mass as a tag mass and use our anchor algorithms for sequencing, addressing the issue of MS data complexity and making 2D HELS MS Seq more robust and accurate.

As it was possible to read segments of up to 35 nt long with a 40K mass resolution LC-MS (*15*), we incorporated a partial RNase T1 digest step to sequence a tRNA^Phe^ that was commercially available, resulting in a reduction of the 76 nt tRNA to fragments of sequenceable sizes. Subsequently, we directly sequenced the entire tRNA with single-base resolution in one single LC-MS run (Fig. 1). To further verify the complete tRNA sequence obtained from the single run above, we labeled the three segments cut by RNase T1 and separated them one by one for 2D-HELS-AA (anchor-based algorithm) MS Seq in three separate LC-MS runs (Fig. S1). To obtain overlapping segment sequences for assembling the complete tRNA, we also included MS data of the tRNA generated without RNase T1 digestion (Fig. 1B). Taking all draft reads output by the anchor-based algorithm together (Table S1-11), we assembled a full length tRNA sequence which was a 100% match to the tRNA^Phe^ reference sequence with more than 2x coverage (Fig.1B). We also successfully identified and located all 11 RNA modifications within the tRNA (Fig. 2). Four of these modifications could be directly read out by their unique masses: dihydrouridine (D) at positions 16 and 17, N^2^, N^2^-dimethylguanosine (m^2^ G) at position 26, 5-methylcytidine (m^5^C) at position 40 and 5-methyluridine (T) at position 54. Methylation on the 2’-OH of C (Cm) at position 32 and G (Gm) at position 34 renders the adjacent 3’-5’-phosphodiester linkage non-hydrolyzable, creating a mass gap in both the 5’- and the 3’-mass ladder families larger than 1 nt (*14*) (Fig. 1A). This gap can be filled by collision induced dissociation (CID) MS, which determines which of the two unhydrolyzable nucleotides is methylated (*14*) (Fig. 2C and Fig. S2). However, other RNA modifications such as pseudouridine (Ψ) and U, N^2^-methylguanosine (m^2^ _2_G) and 7-methylguanosine (m^7^G), and 1-methyladenosine (m^1^A) and N^6^-methyladenosine (m^6^A) share an identical mass, and LC-MS alone cannot distinguish them. Further enzymatic/chemical reactions were required to differentiate m^1^A and m^6^A (*16*), U and Ψ (*15*) (Tables S12-17), and m^2^G and m^7^G (*17*), as shown in the Fig. 2C. Percentages of all the RNA modifications were quantified at each position in case they are not 100% modified (Fig. 2C).

**Fig.1.**
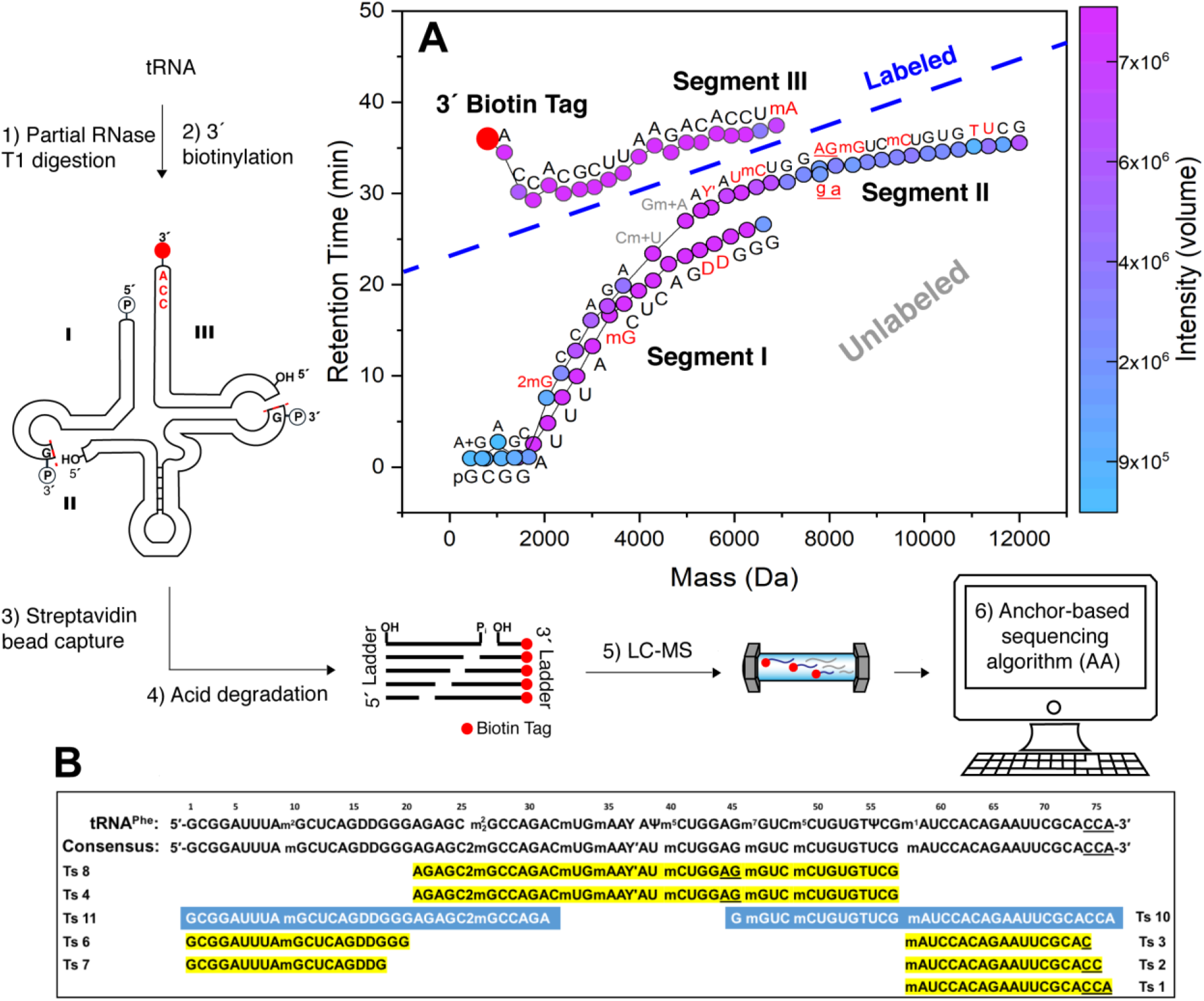
2D-HELS-AA MS Seq of Yeast tRNA^Phe^. **1-6:** Sequencing workflow. More detailed information can be found in Materials and Methods (*supplementary materials*). **A.** The 2D plot of the entire tRNA sequenced from a single LC-MS run, showing the identity and location of all the modifications. **B.** Assembly of the full-length tRNA^Phe^ sequence based on overlapping sequence reads from different LC-MS runs, showing 100% coverage and accuracy as compared to the reported tRNA^Phe^ reference sequence. All output sequence reads are converted to FASTA format in the 5’ to 3’ order (44 and 45 AG conversion output reads not included). *Ts: the Supplementary Table S where the sequencing data of that particular strand can be found.

**Fig. 2.**
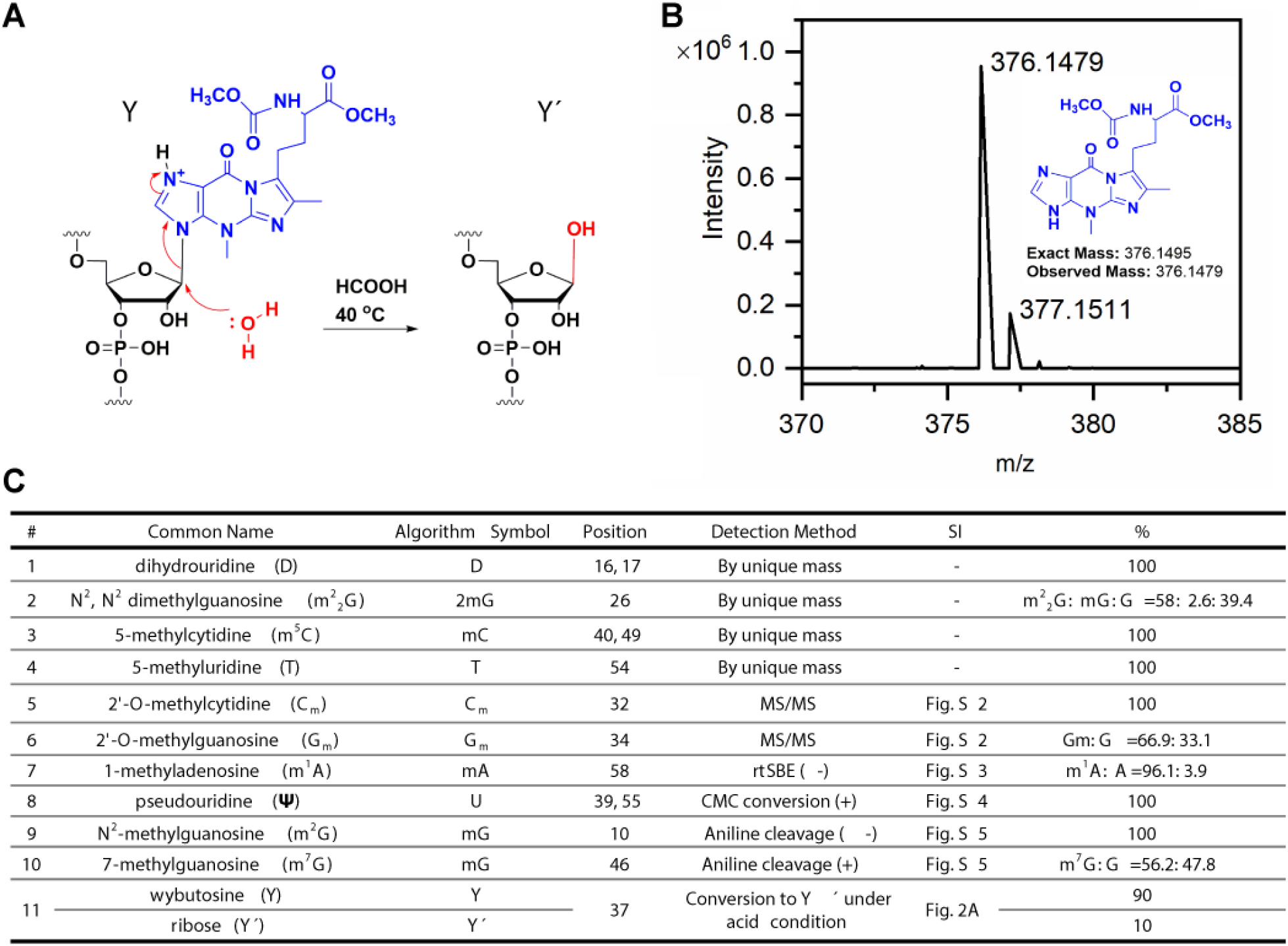
Identification and the subsequent quantification of a dynamic change from wybutosine (Y) to its depurinated Y’ form in tRNA^Phe^. **A.** A proposed mechanism for the conversion of the Y-base to Y’ in acidic conditions. **B.** The mass of nucleobase Y was found in the crude products after acid degradation. The relative percentages of Y and Y’ were quantified and can be found in Table S18. **C.** Summary of all 11 RNA modifications sequenced by 2D-HELS-AA MS Seq. The relative percentages of modifications at each position were quantified by EIC of their corresponding ladder fragments.

Upon analysis of the sequencing results, we noticed that the wybutosine (Y) at position 37 was converted to its depurinated product Y’ (ribose) under acidic degradation conditions (Fig. 2) (*18, 19*). Without acid degradation, only 10% of the tRNA contained the depurinated Y’ form at this position, while 90% contained the standard Y form of the base (Table S18). However, no Y form was observed in any ladder fragments containing this position after acid degradation, and all of the Y bases were converted to Y’ due to depurination with the acidic conditions (Fig. 2A). As another piece of evidence of the depurination, a mass of 376.1178 Da, corresponding to a cleaved Y nucleobase, was found in the crude products after acid degradation and subsequent MS analysis (Fig. 2B), suggesting that Y was originally carried by the tRNA. The fact that our method can identify the dynamic change of Y to Y’ and quantify the relative Y/Y’ ratio could be useful for clinical assays. Certain cancer cells have an acidic pH (*20*), and acid-mediated conversion of Y to Y’ could potentially occur in this pH regime within these cells (*21*). Such changes in the Y’/Y ratio could be used as a potential biomarker for cancer diagnoses. Based on the same principle, our method could potentially probe dynamic changes of other base modifications, and quantify variations in their ratios in particular cells or tissues subjected to different biological processes.

Unlike its nominal identity according to the supplier, upon sequencing, the commercially-prepared tRNA^Phe^ sample was revealed to be heterogeneous. Besides the 76 nt tRNA with a post-transcriptionally modified CCA tail, two other isoforms of the tRNA that are missing an A and a CA at the 3’-CCA tail, respectively, were identified in a 3’ segment of the tRNA (58m^1^A-76A) (Fig.1B and Fig. S6) using the anchor algorithms and a revised Smith-Waterman alignment similarity algorithm (details in *Supplementary Materials*). Surprisingly, the most abundant component was not the nominal 76 nt tRNA^Phe^, which comprised only 17% of the sample as measured by integration of the corresponding extracted-ion currents (EIC) (Fig. S6C) (*11*). Rather the 75 nt tRNA^Phe^ with a missing A at the 3’ end was the major component of the sample, at 80%, while the 74 nt tRNA^Phe^ with a missing CA at the 3’ end was a minor component at 3%. The two tail-truncation isoforms cannot be degraded products of longer tRNAs like the 76 nt tRNA^Phe^, otherwise, they would not have the free 3’-OH required for the 2D HELS chemistry (*15*). The data suggest that 2-D HELS MS Seq is not only is able to sequence modified RNA, it can also identify tail-truncation isoforms that were primarily only studied by polyacrylamide gel electrophoresis methods previously (*22*). As stress-induced tRNA truncation has been implicated in cancers and other diseases (*23*), further studies into the relationship between the relative abundances of tRNA^Phe^ tail-truncation isoforms and various diseases will assist in understanding the role of such isoforms in disease-related biological processes and subsequent treatments (*24*).

Finally, we also observed two isoforms with an A to G transition at position 44 and a G to A transition at position 45, *i.e.*, 44A45G (wild type, reported previously) (*25*) to 44g45a transition (Lower case g and a are used to differentiate them from the canonical G and A in positions 44 and 45, respectively). These two reads were revealed first by our algorithm, and verified manually in the original MFE files (Fig. 3, Tables S4-5, S8-9, and S19-22). We identified two distinct mass ladder fragments at position 44 when reading from the 5’direction, apparently corresponding to sequences containing both 44A and 44g being simultaneously present. However, these two mass ladder fragments merged into one mass ladder fragment at position 45. Such an effect could only occur if two co-existing sequences contained a 45G or a 45a, respectively, thus confirming the coexistence of two co-existing isoforms (Fig. 3A). This is consistent with the sequencing results when reading from the opposite direction when we performed bi-directional sequencing (*14*) (Fig. 3C). We observed these two isoforms in all reads which covered positions 44 and 45, and their relative percentages were consistent (~50% for wild-type, quantified by EIC) (Table S23). To further verify the co-existence of the two mass fragments, we employed full-spectral analysis provided by commercial MassWorks software (Cerno Bioscience, USA) to examine the corresponding ions of these two fragments simultaneously in one spectrum. When reading from the 5’direction, two ions (m/z 778.1051 and 779.7068, both with 10 charge states) were found, corresponding to 44A and 44g. Full-spectral analysis also confirmed that 45G and 45a co-exist when reading from the 3’direction (Fig. 3D). Furthermore, the ratios of 44A/44g as compared to 45G/45a quantified by the full-spectral analysis (*26*) are consistent (Fig. 3), indicating that the sequenced 44g and the 45a are indeed from the same RNA strand, while the 44A and 45G are also both from the same RNA strand. All these MS results support the existence of an isoform, with the sequence 44g45a, co-existing with the wild-type RNA that contains the 44A45G sequence, and that these two isoforms occur at similar levels.

**Fig. 3.**
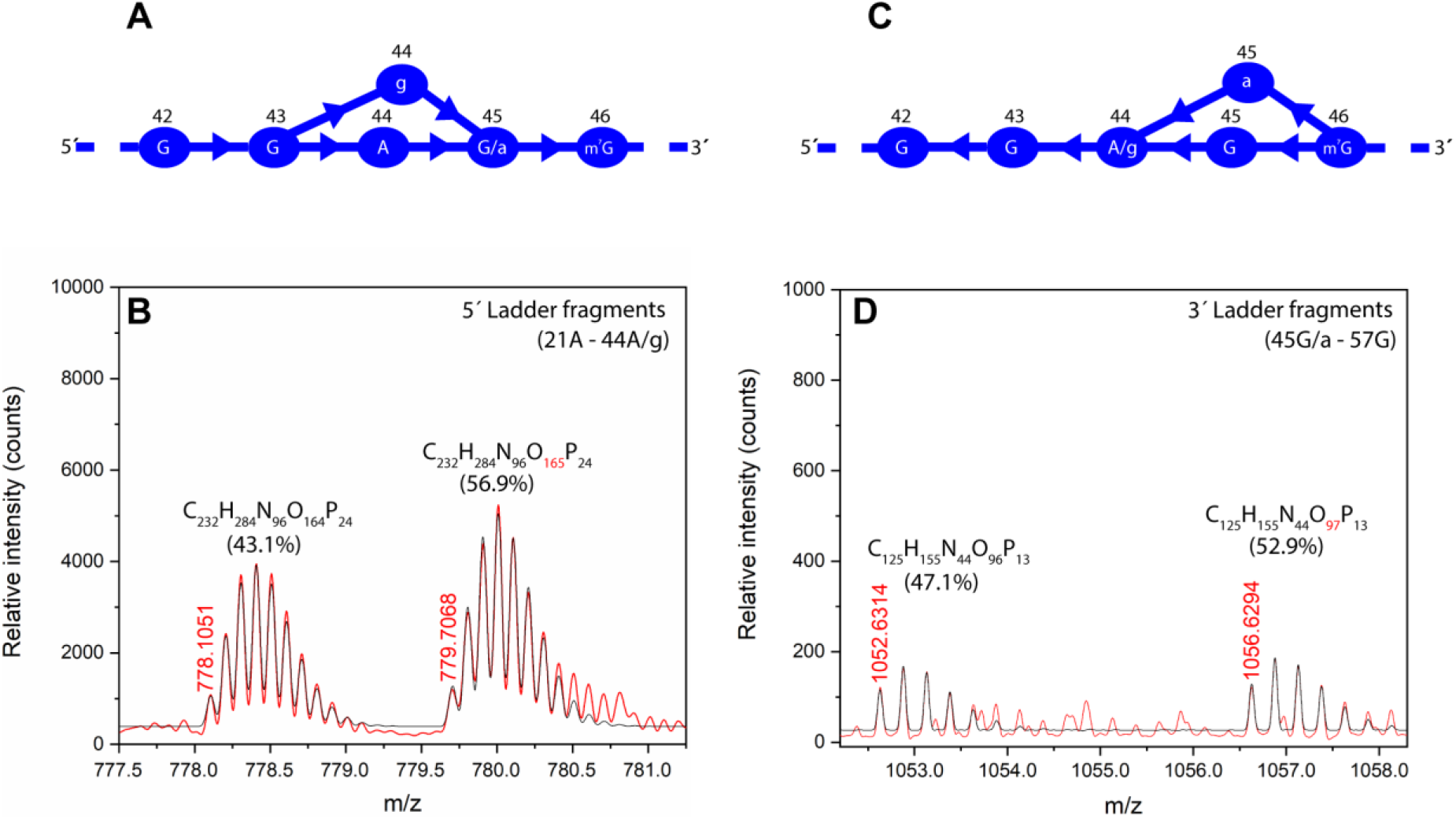
LC-MS results of 44A45G to 44g45a transitions at variable region of tRNA. **A.** A schematic picture of sequence ladder fragments shows a transition g (sharing an identical mass as G) co-exists with A at position 44 when reading from the 5’ direction (Tables S4-5 and S8-9). **B.** Least squares fitted mass spectrum to the calibrated mass spectrum (t_R_ = 31.9-32.9 min) when reading from the 5’ direction. The full-spectral analysis confirms that the ions of the 44g and 44A fragments (with 10 charges) co-exist and that their relative abundances are 57% and 43%, respectively. The theoretical trace (black) of the two combined ion profiles fits well with the calibrated mass spectrum as observed (red), resulting in a good spectral accuracy of 87%. **C.** A single transition a (one oxygen less than G) co-exists with G at position 45 when reading from the 3’ direction (Tables S19-22). **D.** Similarly to **B**, full-spectral analysis confirms that the ions of the 45a and 45G fragments (both with 4 charges; spectral accuracy: 71%) also co-exist and their relative abundances are 47% and 53%, respectively, when reading from the 3’ direction (t_R_ = 16.5-18.6 min).

To further confirm the co-existence of these isoforms, we performed a reverse transcription single base extension (rtSBE) on the tRNA^Phe^ sample. (The rtSBE method was also used in this study to previously differentiate between m^1^A and m^6^A (Fig. S3)). For example, if tRNA^Phe^ has an A/g SNP at position 44, then the rtSBE assay would be able to incorporate both ddT and ddC, since the two isoforms exist at similar levels. However, the results showed that only ddT could be incorporated at position 44 (Fig. S7) and only ddC could be incorporated at position 45 (Fig. S7), indicating that the wildtype 44A45G was the only isoform present. The rtSBE results suggested that RNA reverse transcriptase cannot well recognize these edited bases. It is also possible that the mass differences observed in the above A-G transition at position 44 may be caused by oxidation and reduction, *e.g.*, oxidation of A to isoG and 8-oxoA (Fig. S8A), which both have a mass identical to G. Complete acid digestion of the tRNA into single nucleotides followed by LC-MS analysis support this possibility, as we found two different t_R_s in the EIC profile of the G monophosphate (Fig. S8B), suggesting a co-existing nucleotide of the same mass as G, but a different structure. A similar mechanism could explain the putative G to A transition at position 45. Regardless, the results of the 2D-HELS-AA MS Seq revealed new isoforms, RNA base modifications and editing as well as their relative abundance in the tRNA that can’t be determined by cDNA-based methods (Fig. 4), opening new opportunities in the field of epitranscriptomics.

**Fig. 4.**
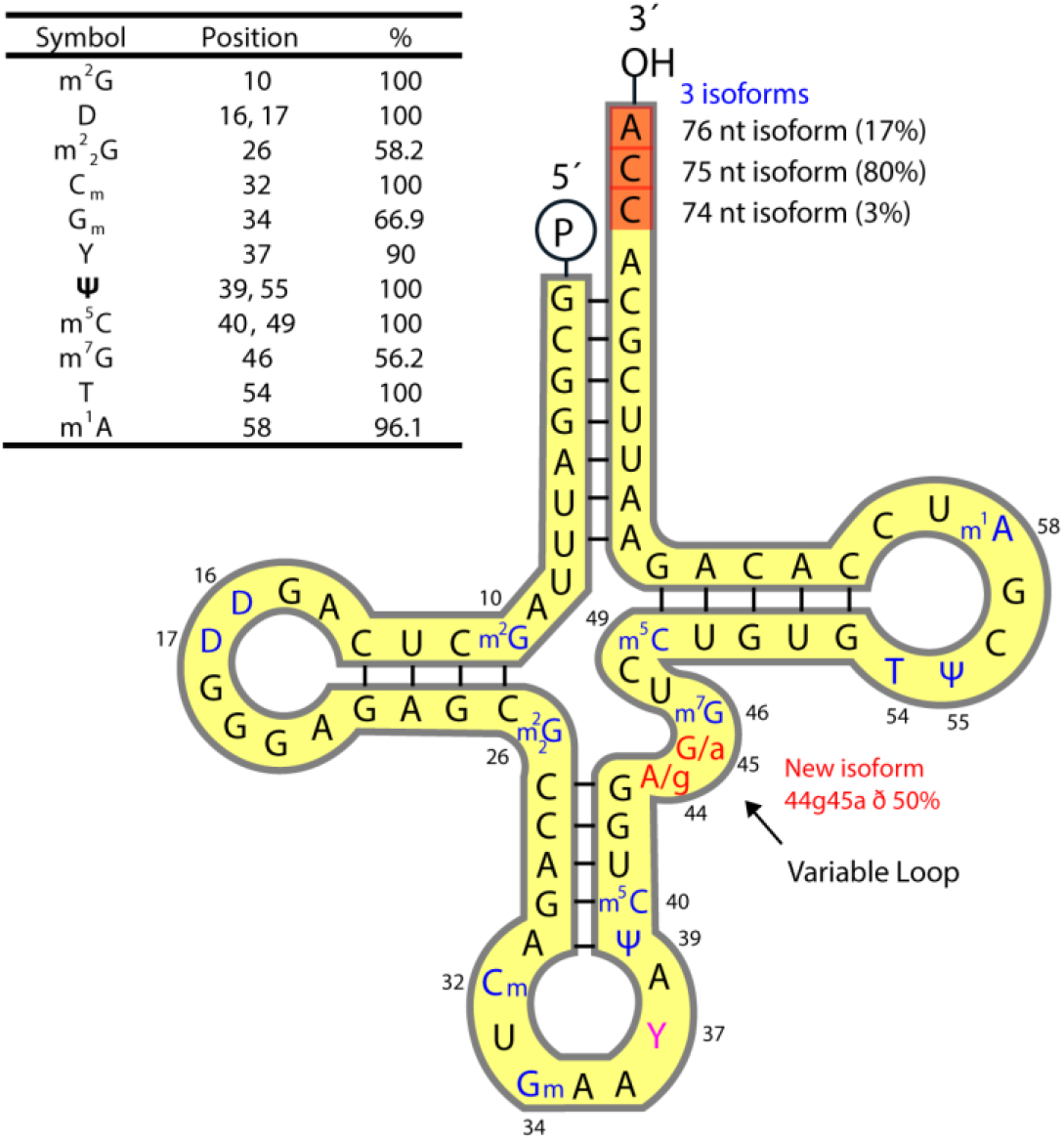
Summary of different isoforms, RNA base modifications and editing as well as their relative abundance in the tRNA^phe^.

## Acknowledgments

The authors acknowledge the R21 grant from NIH (R21HG009576) to S. Z. and W. L. and New York Institute of Technology (NYIT) Institutional Support for Research and Creativity grants to S. Z., which supported this work. The authors would also like to thank PhD student Xuanting Wang (Columbia University) for assisting in figure-making, and Prof. Michael Hadjiargyrou (NYIT), Prof. Jingyue Ju (Columbia University), Drs. Shiv Kumar, Xiaoxu Li, Steffen Jockusch and other members of the Ju lab (Columbia University), Dr. Yongdong Wang (Cerno Bioscience), and Meina Aziz (NYIT) for helpful discussions and suggestions for our manuscript.

## Author contributions

N.Z., S.S., X.W. W.N., T.Z.J., J.J.R., and A.Z. performed experiments and data analysis. B.Y. performed mass spectrometry data acquisition. X.Y., D.J., and W.L. developed anchor-based sequence algorithms, while N.Z., S.S., and S.Z. wrote the manuscript. The original idea for this manuscript was conceived by S.Z.

## Data and materials availability

All data is available in the main text or the supplementary materials. Anchor-based algorithms are available upon request.

