## Supplemental Materials and Methods, Tables and Figs for "2D-HELS-AA MS Seq: Direct sequencing of tRNA reveals its different isoforms and multiple dynamic base modifications"

**This PDF file includes:**

**Materials and Methods**
**Global hierarchical ranking and local best algorithm**
**Isoform detection**
**Figs. S1 - S14**
**Tables S1 - S23**

### Materials and Methods

**Reagent and chemicals:** All chemicals were purchased from commercial sources and used without further purification. tRNA (phenylalanine specific from brewer's yeast), RNaseT1, ATPγS and T4 polynucleotide kinase (3´-phosphatase free) were obtained from Sigma-Aldrich (St. Louis, Missouri, USA), Formic acid (98-100%) was purchased from Merck KGaA (Darmstadt, Germany). Polynucleotide kinase (3´-phosphatase free) and SuperScript IV reverse transcriptase were purchased from Thermo Fisher Scientific (Waltham, MA, USA). Adenosine-5´-5´-diphosphate-{5´-(cytidine-2´-*O*-methyl-3´-phosphate-TEG}-biotin and A(5´)pp(5´)Cp-TEG-biotin-3´ synthesized by ChemGenes (Wilmington, MA, USA). T4 DNA ligase was purchased from New England Biolabs (Ipswich, MA, USA). Biotin maleimide was purchased from Vector Laboratories (Burlingame, CA, USA). All other chemicals, including those needed for conversion of pseudouridine such as CMC (*N*-cyclohexyl-*N′*-(2-morpholinoethyl)-carbodiimide metho-*p*-toluenesulfonate), bicine, urea, EDTA, and $Na_2CO_3$ buffer, were obtained from Sigma-Aldrich unless otherwise stated.

**Dephosphorylation of 5´ end of tRNA:**10 µg of tRNA was digested by 1000 U of RNase T1 followed by purification by Oligo Clean & Concentrator. 20 µL of alkaline phosphatase (20 U/µL, Sigma-Aldrich) was added to the above described tRNA samples and incubated at 50 °C for 60 min followed by purification by Oligo Clean & Concentrator.

**5´ or 3´-ends biotin labeling and biotin streptavidin capture/release purification:** The labeling and purification were performed by previously established methods (*1*).

**tRNA sample preparation for LC-MS (Fig. 1. 1-6):**

**RNase T1 partial digestion and 3´ biotinylation tRNA:** Approximately 10 µg of tRNA was digested by1 µL of 1000 U/µL of RNase T1 in 50 mM Tris-HCl (pH 7.5) containing 2 mM EDTA at room temperature for overnight. The digestion was stopped and purified by Oligo Clean & Concentrator (Zymo Research, Irvine, CA, USA). We monitored the partial digestion by LC-MS and about 40 % tRNA were digested into three fragments, the rest of them were incompletely cut tRNA) and tRNA that was not digested at all. After purification by Oligo Clean & Concentrator, the purified partial digested tRNA were labeled by 3´ biotin (1). After 3´ biotin labeling, we used previously method (*1)* to harvest all 3´ biotinylated RNase T1 partial digested tRNA, which contains 3´ biotin labeled segment III and entire labeled tRNA as well as incompletely cutting fragments. Due to a clover leaf structure of the tRNA, segment I cannot be completely separated from segment III in the un-denatured washing condition used for the biotin-streptavidin capture purification, because segment I has 8 bases that form a stable stem with segment III. The products also include the RNase T1 partial digested tRNA and 3´ biotinlation intact tRNA without any digestion. Indeed, the unlabeled total tRNA and the completely digested segments I and II were mostly removed out during the biotin-streptavidin capture purification and saved for further labeling.

400 µg purified RNase T1 partial digestion and 3´ biotinylation tRNA sample where sequenced by previous method after acid degradation and followed by LC-MS run (*1*).

The acid degraded biotin labeled segment III ladders were shifted up in the 2D plot because $t_R$ shift from the hydrophobic biotin tag. The unlabeled segments I and II ladders were also included in one 2D plot figures but clearly separated from the 3´ biotin labeled segment III ladder. All the sequences were readout by our anchor-based algorithms using the specific anchors.

Labeling Segment II: The residue of purification products were concentrated, desalted by oligo concentrator, and used for next step of 5´ OH biotin labeling segment II. The selective labeling of segment II can be achieved because segment II is the one that has 5´ OH group in the residue and 5´ of the segment I along with the uncut total tRNA have phosphate groups at 5´ ends that cannot be labeled at the same time. The biotin streptavidin capture method was used to purify the 5´ OH biotin labeling of segment II. The residue which contains 5´ of the segment I and uncut total tRNA were saved for the further labeling for the segment I. The labeled segment II was acid degraded for LC-MS sequencing (*1*). The labeled ladder has a $t_R$ shift and can be easily readout by the anchor-based algorithms for further confirming the results from unlabeled segment II in the first run.

Labeling segment I: The materials for labeling segment I can be the residue left from the above segment II labeling step or intact tRNA. Before 5´ OH end labeling step, 5´ dephosphorylation is needed to generate a 5´ OH for labeling 5´ of the segment I and total tRNA, and a similar procedure was used to label segment I.

**LC-MS analysis**

(1) General LC-MS conditions for analyzing tRNA sequencing ladders were the same as previously reported (*1*) except 2-20% buffer B in 60 min followed by a 2 min 90% buffer B wash step.

(2) General MS conditions for the methylated dimers were the same as previously reported.

(3) except the following: targeted ms/ms was used; the mass range for ms1 350-3200 m/z; the mass range for ms2 50-750 m/z. For dimer $C_m$U, the targeted precursor was 642.0837 m/z ($t_R$ = 2.95 min); For dimer $G_m$A, the target precursor was 705.1164 m/z ($t_R$= 3.5 min and 4.08 min), CE = 20. LC conditions: 2-20% MeOH in 60 min (buffer A: 200mM 1,1,1,3,3,3-hexafluoro-2-propanol, 1.25mM triethylamine in water).

(4) General MS conditions for analyzing of single nucleosides or nucleotides if needed were the same as previously reported (*1*) except m/z range 100-2000. LC conditions: 0% B for 5 min, 0-50% B for 30min, 200 µL/min flow; buffer A: water, 0.1% formic acid (FA) and B: acetonitrile (ACN), 0.1% FA, column: Waters Acquity UPLC 2.1×100.

**Computation and data analysis**

The sample data were acquired using the MassHunter Acquisition software (Agilent Technologies, USA). To extract relevant spectral and chromatographic information from the LC-MS experiments, we used the Molecular Feature Extraction (MFE) workflow in MassHunter Qualitative Analysis (Agilent Technologies, USA). This proprietary molecular feature extractor algorithm performs untargeted feature finding in the mass and retention time dimensions. In principal, any software capable of compound identification could be used. The MFE settings were optimized to extract as many identified compounds as possible but with a reasonable quality score. The MFE settings we applied were as follows: "centroid data format, small molecules (chromatographic), peak with height ≥ 100, up to a maximum of 1000, quality score ≥ 30". However, data reduction was performed to simplify algorithm sequencing if needed. For instance, the numbers of input compounds used for algorithm analysis were generally an order-of-magnitude higher than the numbers of ladder fragments needed for generating complete sequences, unless indicated otherwise; these input compounds were sorted out of all MFE extracted compounds typically with higher volumes and/or better quality scores.

The formula used to calculate the PPM in the manuscript: $\mathrm{ppm} = 10^{-6} \times (\mathrm{Mass}_{\mathrm{theoretical}} - \mathrm{Mass}_{\mathrm{observed}})/\mathrm{Mass}_{\mathrm{theoretical}}$

### Global hierarchical ranking and local best algorithm

Data pre-processing is a required step in order for the algorithm to focus on a particular subset of the input dataset at a time. There are two reasons to subset the dataset before parsing into the algorithm. First is to eliminate noise from the dataset. Second is because, experimentally, the RNA material to be sequenced requires fragmentation and labeling with molecular tags. The RNA sample loaded into LC-MS is a mixture of different fragments with some molecular tags. Because of the biochemical properties of the RNA fragments and the tags, in the output dataset from LC-MS, data points corresponding to different RNA fragments are distributed in different groups with distinctive statistics between those groups. The algorithm "zooms in" on one group to read out the sequence of one fragment at a time. Subsetting of the dataset is implemented by refining the RT and mass value of the input dataset in windows, and specifying the starting data point of each fragment. This is feasible because the molecular tag is added to the terminus of each fragment, and the RT and mass feature of the tag is known. Therefore, we call the algorithm "anchor-based", since specifying the starting data point corresponding to the molecular tag latches down the data points corresponding to the specific fragment that we aims to read out from the whole dataset.

After subsetting the dataset, the algorithm performs base calling (Fig. S9). The theoretical mass, calculated from chemical formula, of all known ribonucleotides including those with modifications to the base is stored as a list of $M_{BASE}$. In the first iteration, the algorithm finds the mass corresponding to the molecular tag (anchor) and sets $M_{experimental\_i}$ equal to this mass. The algorithm tests each $M_{BASE}$ from the list by adding it to $M_{experimental\_i}$ and generating a theoretical sum mass $M_{theoretical\_j}$. The algorithm searches through the dataset for a mass value that matches with $M_{theoretical\_j}$. If there exists a matching mass value $M_{experimental\_j}$, a tuple ($M_{experimental\_i}$, *BASE*, $M_{experimental\_j}$) is stored in the result set *V*. Since the algorithm tests all $M_{BASE}$ in the list and looks for all possible matches, multiple tuples with same $M_{experimental\_i}$ but different *BASE* identity and $M_{experimental\_j}$ are stored in set *V*. When the algorithm decides if there is a match, it takes into consideration the experimental error that the experimental mass may slightly deviate from the theoretical mass for a same ribonucleotide. We implemented a calculated parameter PPM that allows $M_{experimental\_j}$ to be matched with $M_{theoretical\_j}$ within a customizable range.

The algorithm performs base calling for all data points until all possible tuples are stored in set *V*. Note that each tuple in set *V* represents an individual base-calling possibility.

After base calling, the algorithm builds trajectories linking tuples in set *V* to generate sequences of the RNA fragment (Fig. S10). Taken tuples from set *V* as vertices, the algorithm finds and stores all edges by examining pairs of tuples such that for a given pair of tuples ($M_i$, *BASE*, $M_j$) and ($M_k$, *BASE*, $M_l$), $M_k = M_j$. The algorithm generates a graph $G = (V, E)$ while finding the edges. When graph *G* is completed, the algorithm finds all paths in graph *G* by depth first search (DFS) (*4*). All paths are stored as sets of vertices. Since the vertices contained in the path are tuples ($M_{experimental\_i}$, *BASE*, $M_{experimental\_j}$), *BASE* can be outputted as a sequence of ribonucleotides.

Because the outputs from LC-MS contains a huge number of data points, graph *G* contains the same number of vertices and also huge number of edges, resulting in tremendous number of total paths, each representing a draft read. To effectively filter the draft reads, we have developed two draft read selection strategies, namely the global hierarchical ranking strategy and the local best score strategy. Nonetheless, both strategies use same parameters acquired from the LC-MS dataset to score the draft reads such as volume and quality score (QS).

In the global hierarchical ranking strategy (Fig. S11 and Fig. S12), the draft reads are scored after the sequence generation step with the following criteria: read length, average volume, average QS, and average PPM. Read length is the number of *BASE* in a draft read. Average volume is calculated by summing the volume associated with each data point in a draft read and diving the sum by read length. Average QS is calculated by dividing the sum of QS by read length for each draft read. Average PPM is the sum of all PPM values associated with data points contained in a draft read divided by read length. The first step of the global hierarchical ranking strategy groups all draft reads into clusters based on their read length, and each cluster is assigned a ranking score for read length. The cluster receiving the highest ranking contains draft reads of the top read length, and the algorithm focuses on this cluster in the following steps. Within this cluster, the draft reads are assigned secondary ranking scores based on average volume values, with drafts reads of higher average volumes receiving higher rankings. In case where more than one draft read have a same read length and average volume value and thus receive a same ranking, the algorithm uses average QS value to re-rank these draft reads, with higher average QS values resulting in higher ranks. If there are still multiple draft reads receiving the same rank, the algorithm uses average PPM value to re-rank these draft reads again, but higher ranks are assigned to draft reads with lower average PPM values since PPM reflects the experimental error associated with each data point from LC-MS. In the end, the draft read with longest read length, highest average volume, highest average QS and lowest average PPM beats all other draft reads in the hierarchical ranking procedure and will be outputted as the final read for the targeted RNA fragment.

Alternatively, the local best score strategy differs from the previous strategy from the step of base calling (Fig. S13 and Fig. S14). The algorithm of local best score strategy applies the anchor-based method to focus on a specific subset of LC-MS dataset presorted by ascending mass order. It pins down the starting ribonucleotide by user defined anchor mass and locates data points from the entire fragment by the anchor. Focusing on these data points, the algorithm now performs base calling and simultaneously evaluates each data point. All data points in the desired zone are now considered as nodes, and the algorithm completes a single path as the final read based on the evaluation of each node. For a current node, it’s mass difference from the previously node (initialized as the anchor) is compared to the list of all known ribonucleotide masses for a match of identity. The match is only accepted if the PPM value of this node is below a certain threshold. In our test data with tRNA samples, we specified this threshold as 10, but it should always be customized to the actual LC-MS dataset. After accepting or rejecting the match (or mismatch otherwise), the algorithm stores the identity of the matched ribonucleotide, and moves on to the next node. There are always several possible next nodes based on their RT. The node with the highest volume will be chosen, with the exception that if a node has outstandingly small PPM value (close to 0) then this node will be chosen over other nodes with higher volumes. The algorithm now searches for a match of identity of the chosen node, evaluates the match, and store the ribonucleotide identity. This process is repeated until the full sequence in the desired data zone is read out.

### CCA truncated isoforms detection

We looked for isoforms of Segment III as an additional step to the global hierarchical ranking algorithm. The final output (Table S1-S3) of the original algorithm is one of the three isoforms and is aligned with all draft reads by Smith-Waterman alignment (*5*) to acquire their alignment score. Draft reads with alignment score above 94.44% are considered candidates of isoforms, and the candidates are ranked by average volume. We acquired six candidates with a cut off at 94.44%. Because the variation between the isoforms is only that they have different tails of C, CC or CCA respectively, the tails of the six candidates were trimmed and a second round of Smith-Waterman alignment was executed. After trimming, draft reads of isoforms had 100% alignment score with each other, and thus were filtered out from the six candidates.

### Chemistry for differentiating pseudouridine (ψ) from uridine

The experiments to convert ψ into CMC-ψ adducts were performed using a modified protocol according to a reported method (*2*). tRNA was denatured in 5 mM EDTA at 80 °C for 2 min and then placed on ice. tRNA (1 nmol) was treated with 0.17 M CMC in 50 mM Bicine (pH 8.3), 4 mM EDTA and 7 M urea at 37 °C for 20 min in a total reaction volume of 90 µL. The reaction was stopped with buffer A (60 µL of 1.5 M sodium acetate and 0.5 mM EDTA, pH 5.6). After purified by Oligo Clean & Concentrator, the resultant product was subsequently treated with 0.05 M $Na_2CO_3$ buffer (pH 10.4) at 37 °C for 17 h. The reaction was stopped with buffer A, and the crude product was purified by Oligo Clean & Concentrator to remove all the salts.

### Chemistry for aniline cleavage at $m^7G$

$tRNA^{Phe}$ (1.6 nmol) was preincubated for 15 min at 37 °C in buffer (Tris-HCl buffer, pH 7.5, 0.01 M $MgCl_2$, 0.2 M KCl). The cooled solution was added to a freshly prepared ice-cold solution of $NaBH_4$ in the same buffer to give final concentrations of 60 µM tRNA and 0.5 M $NaBH_4$. The reduction was performed at 0 °C under subdued light. The reaction was terminated by pipetting aliquots of the reaction mixture into one tenth volume 6 N acetic acid and subsequent purification by Oligo Clean & Concentrator. Then, the tRNA pellet was dissolved in 200 µL × 5 tubes aniline/acetate solution (aniline/acetic acid/water = 1: 3: 7) and incubated for 10 min at 60 °C. 10 volumes of 0.3 M sodium acetate, pH 5.5, were added and subsequently the sample was purified by Oligo Clean & Concentrator.

### Reverse transcription single base extension (rtSBE)

(1) Demethylation: ALKBH3 (2µg/µL) was purchased from Active Motif (CA, USA). The reaction was carried out at 37 °C in 50 mM HEPES buffer (pH 8.0) containing 100 pmol $tRNA^{phe}$, 4µg ALKBH3, 150 µM $Fe(NH_4)_2(SO_4)_2$,1 mM α-ketoglutarate, 2 mM sodium ascorbate, and 1 mM TCEP for 1 h. Oligo Clean & Concentrator was applied to remove salts and excessive reactants.

(2) rtSBE: We designed a reverse primer 5´-TGGTGCGAATTCTGTGGA-3´ (the 3´ primer end is adjacent to the $m^1A$ position), using $tRNA^{phe}$ as a template for $m^1A$ identification, and de-methylated $tRNA^{phe}$ as the control template. The rtSBE reaction was performed by using SuperScript IV reverse transcriptase in 1 × SSIV buffer 30 µl reaction volume containing 25 pmol template, 50 pmol primer, 2.5 nmol ddNTPs, 5 mM DTT, 2 U RNase inhibitor, and 10 U

SuperScript IV reverse transcriptase at 65 °C for 5 min, and then incubated on ice for 1 min. Then reverse transcription reaction was carried out for 25 cycles at 45 °C for 30 sec and 55 °C for 1 min. Lastly, the reaction was inactivated by incubating at 80°C for 10 min followed by using Oligo Clean & Concentrator to remove all salts and proteins. The rtSBE products were measured by MALDI-TOF.

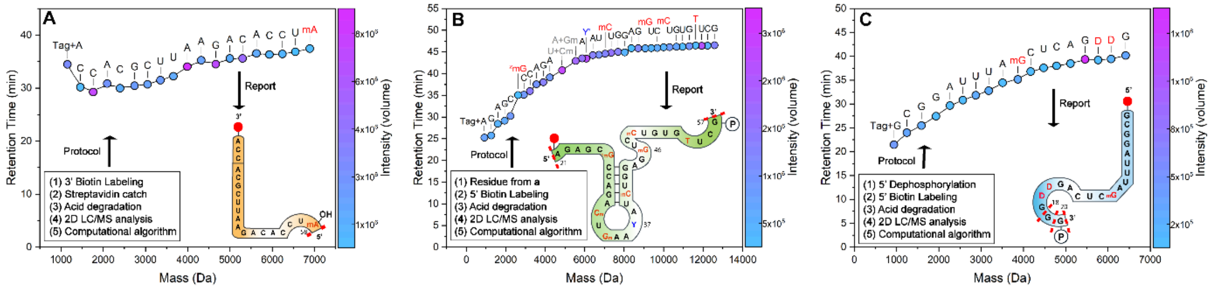

**Fig. S1.** 2D-HELS-AA MS sequencing of three segments digested by T1. As part of HELS, based on the unique chemical moieties in the termini of the three segments, we selectively introduced a biotin label to each of three segments on either the 5´ or 3´ end followed by streptavidin bead-based isolation and release of each segment for acid degradation by formic acid. After LC-MS and data collection, LC-MS data were subsequently exported out by a molecular feature algorithm (MFE, Agilent, USA) for sequence generation using an anchor-based algorithm, which can be implemented either by the global hierarchal ranking or the local best score strategy. We were able to determine a sequence of 19 bases (58m$^1$A to 76A corresponding to segment III (**A**), a sequence of 37 bases (21A to 57G) corresponding to segment II (**B**), and a sequence of 18 bases corresponding to segment I (1G to 18G) (**C**), respectively. We also successfully detected the location of all 11 mass-altering tRNA modifications in the three segments.

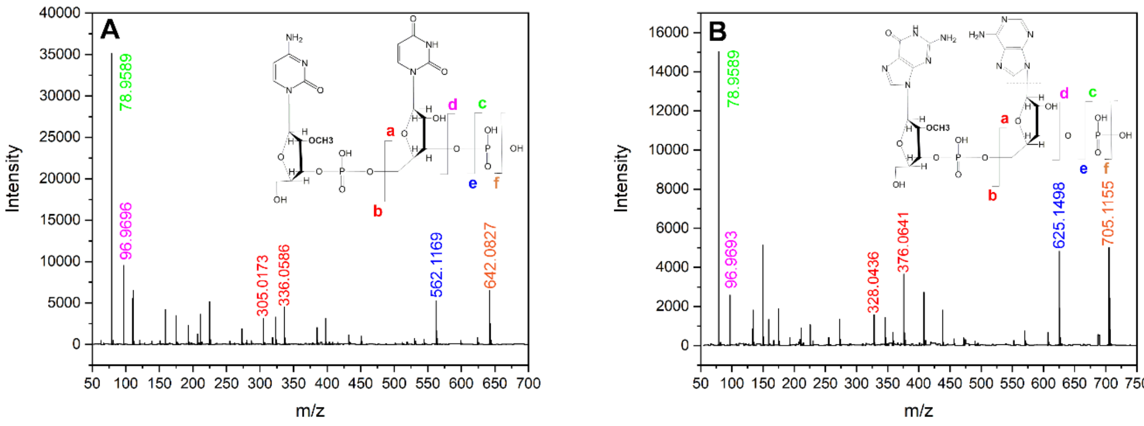

**Fig. S2.** MS analysis of methylated dimers by collision induced dissociation (CID) MS/MS. Samples were prepared by intensive acid hydrolysis (80 °C, 75% (v/v) formic acid, 2h) to generate the dimers. MS/MS data was collected for the modified dimer and fragment ions were used to confirm that the methylation is on the ribose 2´ position of cytidine and the sequence is (**A**) CmU and (**B**) GmA. Assignable fragments labels are indicated on the dimer structures.

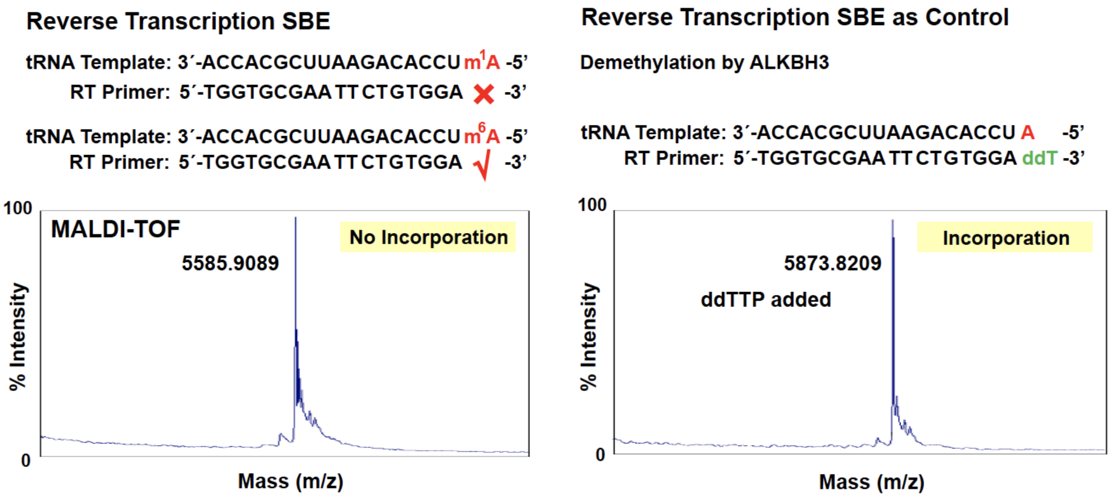


**Fig. S3.** Reverse transcription single base extension (rtSBE) experiment to differentiate m$^1$A and m$^6$A. A pause was observed in the rtSBE experiment, indicating that there is m$^1$A, not m$^6$A, at position 58, because m$^1$A is not able to form base-pairing interactions, thus causing a pause during reverse transcription (*6*).

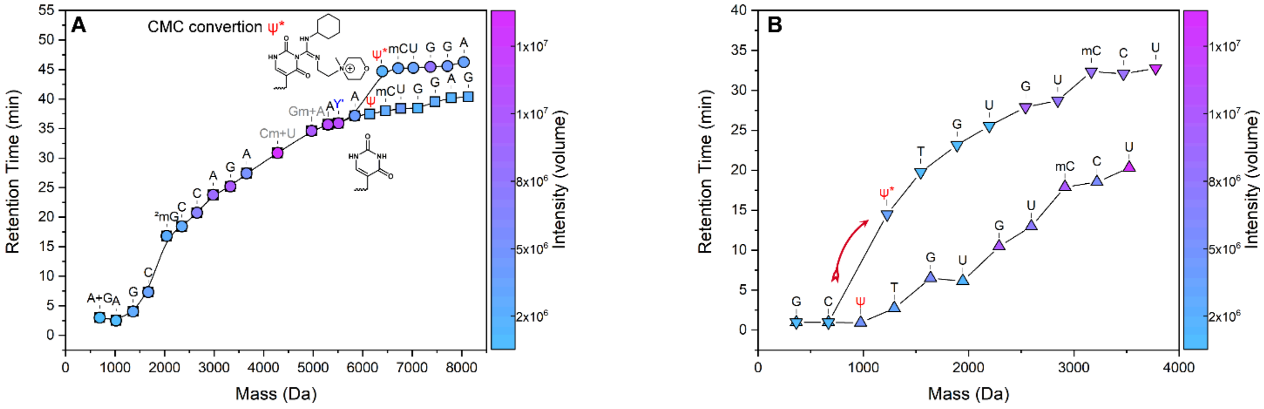


**Fig. S4.** The CMC-converted ψ (depicted as ψ*) results in a shift in both $t_R$ and mass, allowing facile identification and location of ψ at this position due to a single drastic jump in the mass-$t_R$ ladder. For ease of visualization, only the sequences of the **(A)** 5´-mass-tR ladder (22G to 44A) and **(B)** 3´-mass-$t_R$ ladder (57G to 47U) are presented. The sequences presented were manually acquired based on the mass-$t_R$ ladders identified from the algorithm-processed data. Insert picture in **(A)** shows the chemical conversion of ψ by reacted with CMC to form the CMC-ψ adduct, shifting CMC-ψ-containing mass-$t_R$ ladders in both mass and $t_R$ compared to mass-$t_R$ ladders containing unconverted ψ.

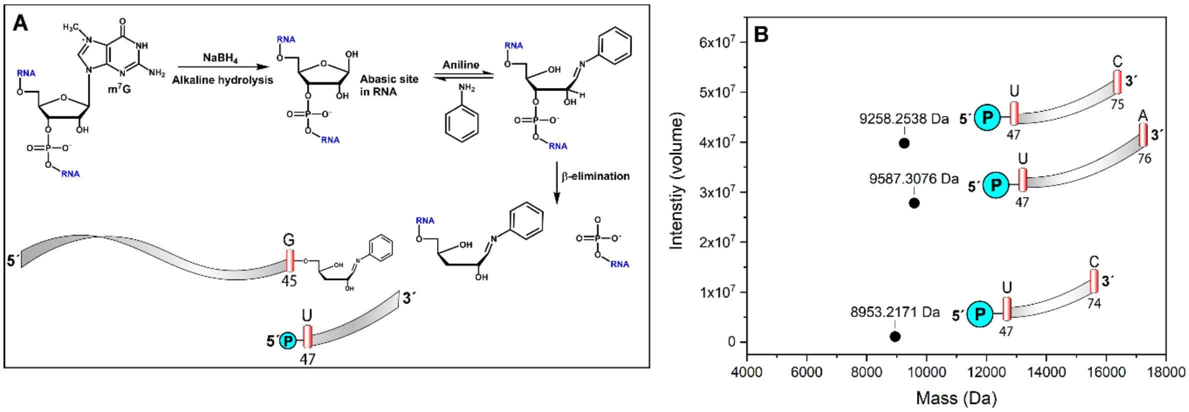


C

| Fragment | Calc mass | Exp mass | m/z | EIC (Area) | Quality score | ppm |
|---|---|---|---|---|---|---|
| 47U to 75C | 9258.2251 | 9258.2538 | 711.1664 | 11191706 | 100 | -3.1 |
| 47U to 76A | 9587.2776 | 9587.3076 | 736.4783 | 9197206 | 100 | -3.1 |

**Fig. S5. (A)** Chemistry for distinguishing $m^7G$ from other isomer base modifications such as $m^2G$ that share an identical mass. **(B)** The plot of Mass *vs.* Intensity after the chemical cleavage of the RNA at $m^7G$ site-specifically. The mass of the three major fragments observed were 9587.3076 Da, 9258.2538 Da and 8953.2171 Da, corresponding to their 76 nt, 75 nt and 74 nt isoforms, respectively, indicating that there is a $m^7G$ at the 46 position. **(C)** Specific fragments cleaved at $m^7G$ were analyzed by LC-MS and quantified by integrating the extracted ion current (EIC).

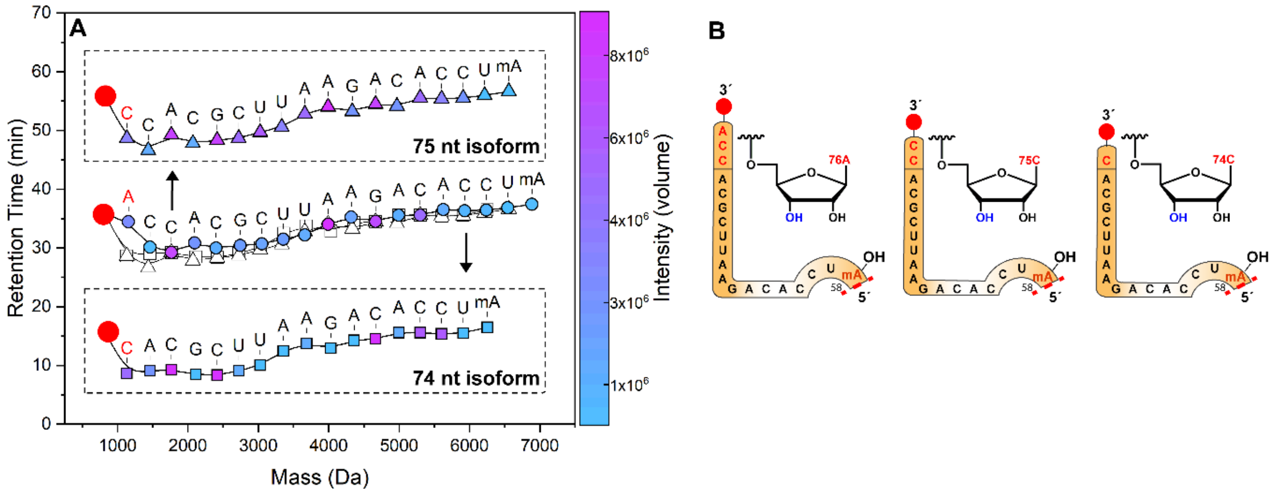

**C**

| Fragment | Calc mass | Exp mass | m/z | EIC (Area) | Percent | Quality score | ppm |
|---|---|---|---|---|---|---|---|
| 58 $m^1A$ to 74C | 5364.7935 | 5364.7939 | 595.0800 | 226450 | 3% | 98 | -0.1 |
| 58 $m^1A$ to 75C | 5669.8348 | 5669.8403 | 628.9753 | 6242830 | 80% | 100 | -1.0 |
| 58 $m^1A$ to 76A | 5998.8873 | 5998.8845 | 598.8828 | 1323018 | 17% | 100 | 0.5 |

**Fig. S6. (A)** Sequencing of Segment III and identification of two other isoforms of $tRNA^{Phe}$ with a terminal 75C and 74C at the 3´ end. $t_R$ was normalized for ease of visualization. LC-MS analysis of 3´-biotin-labeled mass ladders of tRNA after RNase T1 digestion followed by subsequent chemical degradation. The sequences presented were manually acquired based on the mass-$t_R$ ladders identified from the algorithm-processed data. The insets represent the two truncated isoforms separated from the rest of the data for ease of visualization. (**B**) The terminal base of 76 nt $tRNA^{Phe}$, and its two tail-truncated isoforms; all three isoforms contain a free OH at the 3´ for introducing the biotin tag, suggesting that the isoforms were not generated during acid degradation. (C) RNase T1 digestion products of tRNA measured by LC-MS. Among them, three major segments were observed which have the strongest peak volume. The relative quantities of different product species were quantified by EIC (*1*, *7*).

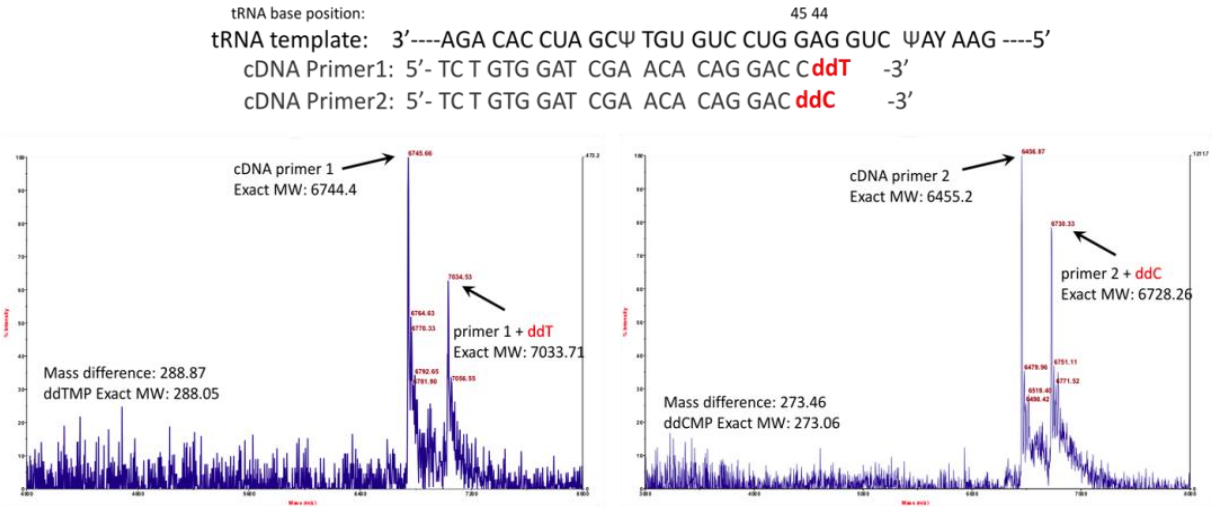


**Fig. S7.** MALDI-TOF results of reverse transcription Single Base Extension (rtSBE): for cDNA primer1, only ddT(44) was incorporated and for cDNA primer2, only ddC(45) was incorporated. The results supported the tRNA template was 44A and 45 G wild type isoform.

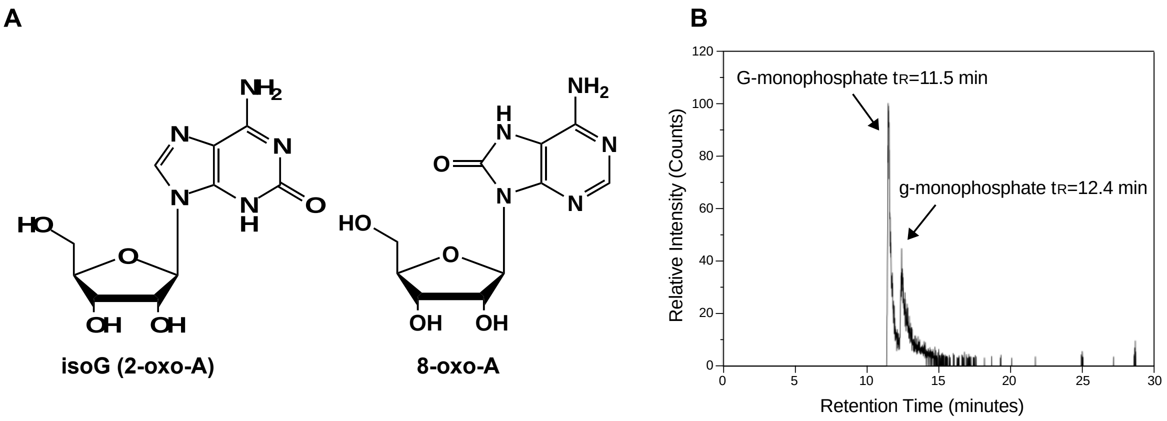


**Fig. S8.** (A) Chemical structures of isoG and 8-oxo-A. (B) we found two different $t_R$s in the EIC profile of the G monophosphate, suggesting a co-existing nucleotide of the same mass as G.

**Base-calling**

Input: Experimentally acquired LC-MS dataset of sample RNA (dataset)
Theoretical mass of molecular tag label (anchor)

Output: Result set $V = (M_{experimental\_i}, BASE, M_{experimental\_j})$ ,where $M_{experimental\_i}$ and $M_{experimental\_i}$ are mass values found in the dataset and *BASE* is the identity of the ribonucleotide detected in the base calling

FOR each mass value $M_{experimental\_i}$ $(i = 0, 1, 2,\ldots, n)$ in dataset
    FOR each mass value $M_{experimental\_j}$ $(j > i)$ in dataset
        FOR each $M_{BASE}$
            $M_{theoretical\_j} = M_{experimental\_i} + M_{BASE}$
            IF $| M_{theoretical\_j} - M_{experimental\_j} | <$ PPM
                Add matched tuple $(M_{experimental\_i}, BASE, M_{experimental\_j})$ to set $V$
            END IF
        END FOR
    END FOR
END FOR

**Fig. S9.** The pseudocode for base-calling step of the global hierarchical ranking algorithm. In this step the algorithm stores all possible tuples of $(M_i, BASE, M_j)$ recoding the mass from MS data as $M_i$ and $M_j$ and the base identity matching with the mass difference of $M_i$ and $M_j$ as *BASE*.

**Sequence generation**

Input: Vertex set $V = (M_{experimental\_i}, BASE, M_{experimental\_j})$

Output: Ribonucleotide trajectories (sequences)

```
FOR each vertex A (M_i, BASE, M_j) in set V
        FOR each vertex B (M_k, BASE, M_l) in set V
                IF M_k = M_j
                        Add edge e = (A, B) to edge set E
                END IF
        END FOR
END FOR

Find all paths in graph G = (V, E) by depth first search (DFS)
A path is composed of multiple edges from edge set E connecting a set of vertices (v_1, v_2, …, v_n)

Output the paths (BASE_1, BASE_2, BASE_3,… )
```

**Fig. S10.** The pseudocode for sequence generation step of the global hierarchical ranking algorithm. In this step the algorithm takes the tuples stored in base-calling as nodes and connects the nodes to build paths corresponding to draft reads.

**Draft reads selection**

Input: All paths in graph *G*
Volume, quality score (QS), PPM from LC-MS dataset

Output: Final read of RNA sequence

```
FOR each path in graph G
        Calculate read length
        Calculate average volume, average QS, average PPM
END FOR

Cluster all paths by read length
        Rank clusters

In the top ranking cluster for read length
        Rank draft reads by average volume
                Rank draft reads by average QS
                        Rank draft reads by average PPM

Output final read
```

**Fig. S11.** The pseudocode of the draft read selection step of the global hierarchical ranking algorithm. The draft reads are evaluated by four parameters in order: read length, average volume, average QS and average PPM, which each parameter the algorithm performs a round of ranking of the draft reads. The draft read at the top ranking becomes the final output.

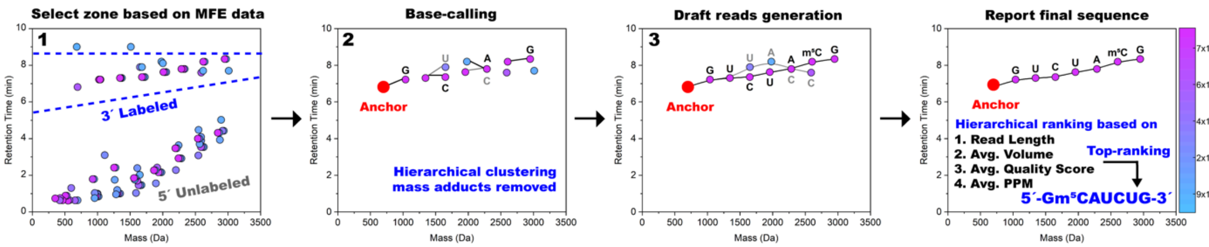


**Fig. S12**. Workflow of data analysis using an anchor-based sequencing algorithm with the global hierarchical ranking strategy. The MS data shown in the workflow is simulated with a purified sample, and the intensity of the color indicates the associated volume of each data point with darker blue points indicating higher volume and *vice versa*. $Na^+$, $2Na^+$, $Na^+$+$K^+$ and other mass adducts were hierarchically clustered to augment compound intensity and to reduce data complexity in step 2. The processed data were subsetted by filtering $t_R$ and mass value, so that only data points in the zone of labeled fragments were passed on in the algorithm in step 1. An anchor-based algorithm was applied for *de novo* sequence generation automatically. All draft reads were ranked by read length, average volume, average QS and average PPM in this order, and the top-ranking draft read for each fragment was output and chosen as the final output read.

```
Best local score algorithm
Input:          Experimentally acquired LC-MS dataset of sample RNA (dataset)
                Theoretical mass of molecular tag (M_anchor)

Output:         Path of final read P = (BASE_1, BASE_2, BASE_3, ......)

Sort dataset by order of ascending mass value
M_i = M_anchor
WHILE  M_k  in dataset
        IF M_K in range (mass_lower_bound, mass_upper_bound)
                IF RT_k in range (RT_lower_bound, RT_upper_bound)
                        IF PPM_k < threshold
                                M_gap = M_k – M_i
                                FOR each M_BASE
                                        IF M_BASE = M_gap
                                                Add BASE to path P
                                        END IF
                                END FOR
                        END IF
                END IF
        END IF
        M_i = M_k
        M_k = next node whose PPM < 0.11 (extremely low value, close to 0); or next node with highest volume
END WHILE
```

**Fig. S13.** The pseudocode of the local best score algorithm. Instead of generating all possible tuples during base calling, the local best score algorithm only stores the base identity and corresponding mass with the highest volume. Thus, the local best score algorithm generates only one draft read.

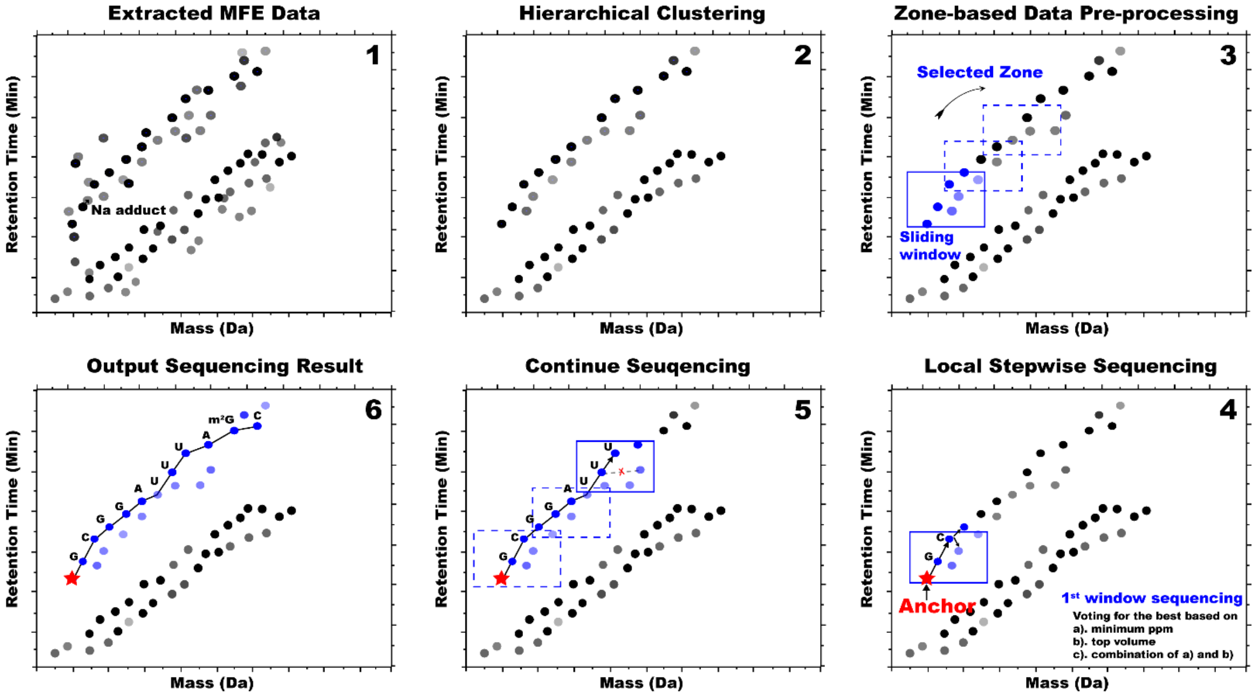


**Fig. S14.** The algorithm implementing the local best score strategy, performed by a Python coding system.

**Table S1.** 3´_biotin_tRNA_T1_SIII_111418s05_76A. Sequencing of 3´ biotin labeled tRNA segment III from 58m$^1$A to 76A by global hierarchical ranking algorithm.

| Fragment | Mass | RT | Base | Volume | PPM |
|---|---|---|---|---|---|
| 1 | 826.3164 | 35.809 | Tag | 2645323 | 2.42 |
| 2 | 1155.3679 | 34.555 | A | 580850 | 2.60 |
| 3 | 1460.4116 | 30.202 | C | 259583 | 0.41 |
| 4 | 1765.4505 | 29.311 | C | 4875476 | 1.70 |
| 5 | 2094.5027 | 30.921 | A | 560348 | 1.58 |
| 6 | 2399.5455 | 30.024 | C | 241970 | 0.75 |
| 7 | 2744.5948 | 30.494 | G | 365785 | 0.04 |
| 8 | 3049.6138 | 30.755 | C | 245795 | 7.28 |
| 9 | 3355.6561 | 31.570 | U | 377273 | 1.58 |
| 10 | 3661.6854 | 32.930 | U | 4226311 | 0.38 |
| 11 | 3990.7364 | 34.122 | A | 4968527 | 0.73 |
| 12 | 4319.7918 | 35.332 | A | 245329 | 0.00 |
| 13 | 4664.8388 | 34.606 | G | 4756748 | 0.09 |
| 14 | 4993.8992 | 35.504 | A | 307359 | 1.50 |
| 15 | 5298.9333 | 35.691 | C | 4083332 | 0.06 |
| 16 | 5627.9522 | 35.501 | A | 160811 | 5.92 |
| 17 | 5933.0022 | 35.649 | C | 157328 | 4.15 |
| 18 | 6238.0838 | 36.541 | C | 89737 | 2.52 |
| 19 | 6544.1101 | 36.202 | U | 672814 | 2.54 |
| 20 | 6887.1727 | 37.539 | mA | 1193510 | 1.61 |

>Ts1
mAUCCACAGAAUUCGCACCA

**Table S2.** 3´_biotin_tRNA_T1_SIII_111418s05_75C. Sequencing of 3´ biotin labeled tRNA segment III from 58m$^1$A to 75C by global hierarchical ranking algorithm.

| Fragment | Mass | RT | Base | Volume | PPM |
|---|---|---|---|---|---|
| 1 | 826.3164 | 35.809 | Tag | 2645323 | 2.42 |
| 2 | 1131.3573 | 28.724 | C | 2536602 | 2.12 |
| 3 | 1436.3979 | 26.748 | C | 1504369 | 2.16 |
| 4 | 1765.4505 | 29.311 | A | 4875476 | 1.70 |
| 5 | 2070.4898 | 27.904 | C | 1807879 | 2.41 |
| 6 | 2415.5392 | 28.436 | G | 4919858 | 1.24 |
| 7 | 2720.5806 | 28.781 | C | 4403013 | 1.07 |
| 8 | 3026.6061 | 29.745 | U | 5263366 | 0.93 |
| 9 | 3332.6311 | 30.654 | U | 3654432 | 0.96 |
| 10 | 3661.6854 | 32.930 | A | 4226311 | 0.38 |
| 11 | 3990.7364 | 34.122 | A | 4968527 | 0.73 |
| 12 | 4335.7879 | 33.348 | G | 2855812 | 0.28 |
| 13 | 4664.8388 | 34.606 | A | 4756748 | 0.09 |
| 14 | 4969.8783 | 34.250 | C | 2303352 | 0.44 |
| 15 | 5298.9333 | 35.691 | A | 4083332 | 0.06 |
| 16 | 5603.9769 | 35.502 | C | 2292626 | 0.46 |
| 17 | 5909.0178 | 35.637 | C | 2429322 | 0.37 |
| 18 | 6215.0412 | 36.088 | U | 860704 | 0.03 |
| 19 | 6558.1157 | 36.751 | mA | 16787962 | 1.01 |

>Ts2
mAUCCACAGAAUUCGCACC

**Table S3.** 3´_biotin_tRNA_T1_SIII_111418s05_74C. Sequencing of 3´ biotin labeled tRNA segment III from 58m$^1$A to 74C by global hierarchical ranking algorithm.

| Fragment | Mass | RT | Base | Volume | PPM |
|---|---|---|---|---|---|
| 1 | 826.3164 | 35.809 | Tag | 2645323 | 2.42 |
| 2 | 1131.3573 | 28.724 | C | 2536602 | 2.12 |
| 3 | 1460.4116 | 30.202 | A | 259583 | 0.41 |
| 4 | 1765.4505 | 29.311 | C | 4875476 | 1.70 |
| 5 | 2110.4918 | 27.882 | G | 356221 | 4.31 |
| 6 | 2415.5392 | 28.436 | C | 4919858 | 1.24 |
| 7 | 2721.5695 | 29.145 | U | 239635 | 0.70 |
| 8 | 3027.5972 | 30.047 | U | 68400 | 1.39 |
| 9 | 3356.6432 | 32.543 | A | 189932 | 0.69 |
| 10 | 3685.6934 | 33.833 | A | 159564 | 1.25 |
| 11 | 4030.7417 | 33.004 | G | 82558 | 0.92 |
| 12 | 4359.8007 | 34.352 | A | 289735 | 0.64 |
| 13 | 4664.8388 | 34.606 | C | 4756748 | 0.09 |
| 14 | 4993.8992 | 35.504 | A | 307359 | 1.50 |
| 15 | 5298.9333 | 35.691 | C | 4083332 | 0.06 |
| 16 | 5603.9769 | 35.502 | C | 2292626 | 0.46 |
| 17 | 5910.0206 | 35.639 | U | 98526 | 3.54 |
| 18 | 6253.0697 | 36.605 | mA | 181155 | 0.30 |

>Ts3
mAUCCACAGAAUUCGCAC

**Table S4.** 5´_OH_tRNA_T1_SII_111418s05_44A45G. Sequencing of 5´ OH tRNA segment II from 21A to 57G by global hierarchical ranking algorithm.

| Fragment | Mass | RT | Base | Volume | PPM |
|---|---|---|---|---|---|
| 1 | 692.1081 | 0.945 | A+G | 448392 | 3.47 |
| 2 | 1021.1592 | 0.996 | A | 612623 | 3.72 |
| 3 | 1366.2059 | 1.023 | G | 1163701 | 3.29 |
| 4 | 1671.2489 | 1.112 | C | 1917190 | 1.68 |
| 5 | 2044.3269 | 8.858 | 2mG | 2025885 | 1.71 |
| 6 | 2349.3682 | 10.309 | C | 3120462 | 1.49 |
| 7 | 2654.4101 | 12.749 | C | 6309574 | 1.09 |
| 8 | 2983.4617 | 16.073 | A | 5462129 | 1.27 |
| 9 | 3328.5102 | 17.647 | G | 6892234 | 0.81 |
| 10 | 3657.5632 | 19.875 | A | 4203490 | 0.60 |
| 11 | 4282.6476 | 23.391 | U+Cm | 11059167 | 0.02 |
| 12 | 4970.7632 | 26.996 | A+Gm | 8957192 | 0.02 |
| 13 | 5299.8175 | 28.115 | A | 9137581 | 0.32 |
| 14 | 5511.8281 | 28.449 | Y' | 9044373 | 0.67 |
| 15 | 5840.8796 | 29.718 | A | 7213450 | 0.46 |
| 16 | 6146.9082 | 30.061 | U | 12938074 | 0.98 |
| 17 | 6465.9647 | 30.688 | mC | 6445803 | 0.87 |
| 18 | 6771.9918 | 31.161 | U | 6802824 | 1.09 |
| 19 | 7117.0401 | 31.251 | G | 3468612 | 1.17 |
| 20 | 7462.0865 | 32.049 | G | 2834683 | 0.98 |
| 21 | 7791.1394 | 32.735 | A | 2239278 | 1.00 |
| 22 | 8136.1981 | 33.016 | G | 3437631 | 2.35 |
| 23 | 8495.2645 | 33.131 | mG | 2251492 | 2.62 |
| 24 | 8801.2888 | 33.439 | U | 3178250 | 2.42 |
| 25 | 9106.3319 | 33.677 | C | 3146668 | 2.54 |
| 26 | 9425.3892 | 33.961 | mC | 3341188 | 2.49 |
| 27 | 9731.4100 | 34.135 | U | 3700286 | 1.95 |
| 28 | 10076.4607 | 34.378 | G | 2776140 | 2.21 |
| 29 | 10382.4798 | 34.582 | U | 2849708 | 1.55 |
| 30 | 10727.5480 | 34.793 | G | 2740634 | 3.44 |
| 31 | 11047.5761 | 35.136 | T | 781981 | 2.17 |
| 32 | 11353.6241 | 35.183 | U | 4303300 | 4.11 |
| 33 | 11658.6776 | 35.364 | C | 1498752 | 5.05 |
| 34 | 12003.6973 | 35.531 | G | 6123452 | 2.60 |

>Ts4
AGAGC2mGCCAGACmUGmAAY'AUmCUGGAGmGUCmCUGUGTUCG

**Table S5.** 5´_OH_tRNA_T1_SII_111418s05_44g45a. Sequencing of 5´ OH tRNA segment II from 21A to 57G by global hierarchical ranking algorithm.

| Fragment | Mass | RT | Base | Volume | PPM |
|---|---|---|---|---|---|
| 1 | 692.1081 | 0.945 | A+G | 448392 | 3.47 |
| 2 | 1021.1592 | 0.996 | A | 612623 | 3.72 |
| 3 | 1366.2059 | 1.023 | G | 1163701 | 3.29 |
| 4 | 1671.2489 | 1.112 | C | 1917190 | 1.68 |
| 5 | 2044.3269 | 8.858 | 2mG | 2025885 | 1.71 |
| 6 | 2349.3682 | 10.309 | C | 3120462 | 1.49 |
| 7 | 2654.4101 | 12.749 | C | 6309574 | 1.09 |
| 8 | 2983.4617 | 16.073 | A | 5462129 | 1.27 |
| 9 | 3328.5102 | 17.647 | G | 6892234 | 2.55 |
| 10 | 3657.5632 | 19.875 | A | 4203490 | 0.60 |
| 11 | 4282.6476 | 23.391 | U+Cm | 11059167 | 0.02 |
| 12 | 4970.7632 | 26.996 | A+Gm | 8957192 | 0.02 |
| 13 | 5299.8175 | 28.115 | A | 9137581 | 0.34 |
| 14 | 5511.8281 | 28.449 | Y' | 9044373 | 0.69 |
| 15 | 5840.8796 | 29.718 | A | 7213450 | 0.48 |
| 16 | 6146.9082 | 30.061 | U | 12938074 | 0.98 |
| 17 | 6465.9647 | 30.688 | mC | 6445803 | 0.87 |
| 18 | 6771.9918 | 31.161 | U | 6802824 | 1.08 |
| 19 | 7117.0401 | 31.251 | G | 3468612 | 1.15 |
| 20 | 7462.0865 | 32.049 | G | 2834683 | 0.96 |
| 21 | 7807.1332 | 32.101 | G | 2248564 | 0.83 |
| 22 | 8136.1981 | 33.016 | A | 3437631 | 2.32 |
| 23 | 8495.2645 | 33.131 | mG | 2251492 | 2.61 |
| 24 | 8801.2888 | 33.439 | U | 3178250 | 2.40 |
| 25 | 9106.3319 | 33.677 | C | 3146668 | 2.51 |
| 26 | 9425.3892 | 33.961 | mC | 3341188 | 2.47 |
| 27 | 9731.4100 | 34.135 | U | 3700286 | 1.92 |
| 28 | 10076.4607 | 34.378 | G | 2776140 | 2.18 |
| 29 | 10382.4798 | 34.582 | U | 2849708 | 1.51 |
| 30 | 10727.5480 | 34.793 | G | 2740634 | 3.40 |
| 31 | 11047.5761 | 35.136 | T | 781981 | 2.14 |
| 32 | 11353.6241 | 35.183 | U | 4303300 | 4.07 |
| 33 | 11658.6776 | 35.364 | C | 1498752 | 5.01 |
| 34 | 12003.6973 | 35.531 | G | 6123452 | 2.56 |

>Ts5
AGAGC2mGCCAGACmUGmAAY'AUmCUGGGAmGUCmCUGUGTUCG

**Table S6.** 5´_pG_tRNA_T1_SI_111418s05. Sequencing of 5´ pG tRNA segment I from 1G to 20G by global hierarchical ranking algorithm.

| Fragment | Mass | RT | Base | Volume | PPM |
|---|---|---|---|---|---|
| 1 | 443.0222 | 0.968 | pG | 32204 | 4.74 |
| 2 | 748.0626 | 0.935 | C | 327973 | 4.01 |
| 3 | 1093.1092 | 0.963 | G | 247078 | 3.48 |
| 4 | 1438.1583 | 1.010 | G | 1953624 | 1.46 |
| 5 | 1767.2105 | 2.512 | A | 6646248 | 1.36 |
| 6 | 2073.2377 | 4.800 | U | 11078570 | 0.24 |
| 7 | 2379.2611 | 7.664 | U | 13653044 | 1.01 |
| 8 | 2685.2874 | 9.948 | U | 13651928 | 0.52 |
| 9 | 3014.3399 | 13.244 | A | 8446589 | 0.46 |
| 10 | 3373.3974 | 16.657 | mG | 5400820 | 2.08 |
| 11 | 3678.4462 | 17.883 | C | 6427287 | 0.14 |
| 12 | 3984.4711 | 19.330 | U | 10498687 | 0.03 |
| 13 | 4289.5141 | 20.432 | C | 13067020 | 0.42 |
| 14 | 4618.5661 | 22.240 | A | 9336602 | 0.28 |
| 15 | 4963.6167 | 23.110 | G | 19445698 | 0.91 |
| 16 | 5271.6368 | 23.792 | D | 6241383 | 3.11 |
| 17 | 5579.6992 | 24.454 | D | 7740033 | 0.90 |
| 18 | 5924.7535 | 25.268 | G | 104745696 | 2.01 |
| 19 | 6269.8003 | 25.980 | G | 3057757 | 1.80 |
| 20 | 6614.8364 | 26.615 | G | 673220 | 0.00 |

>Ts6
GCGGAUUUAmGCUCAGDDGGG

**Table S7.** 5´_biotin_tRNA_T1_SI_042519s07. Sequencing of 5´ biotin labeled tRNA segment I from 1G to 18G by global hierarchical ranking algorithm.

| Fragment | Mass | RT | Base | Volume | PPM |
|---|---|---|---|---|---|
| 1 | 938.2184 | 21.449 | Tag+G | 403806 | 3.41 |
| 2 | 1243.2600 | 23.971 | C | 277726 | 2.33 |
| 3 | 1588.3060 | 25.493 | G | 238503 | 2.71 |
| 4 | 1933.3518 | 27.433 | G | 44902 | 3.05 |
| 5 | 2262.4042 | 29.682 | A | 35264 | 2.65 |
| 6 | 2568.4387 | 30.807 | U | 64428 | 1.21 |
| 7 | 2874.4631 | 31.835 | U | 219666 | 0.73 |
| 8 | 3180.4871 | 32.783 | U | 173234 | 0.22 |
| 9 | 3509.5467 | 34.465 | A | 67573 | 2.22 |
| 10 | 3868.6148 | 35.174 | mG | 226704 | 3.31 |
| 11 | 4173.6443 | 36.794 | C | 63409 | 0.24 |
| 12 | 4479.6520 | 37.559 | U | 12772 | 3.73 |
| 13 | 4784.7078 | 38.002 | C | 14478 | 0.46 |
| 14 | 5113.7758 | 38.479 | A | 69348 | 2.60 |
| 15 | 5458.8177 | 39.347 | G | 1588901 | 1.43 |
| 16 | 5766.8095 | 39.208 | D | 25595 | 7.18 |
| 17 | 6074.9000 | 39.440 | D | 118414 | 1.33 |
| 18 | 6419.9573 | 40.140 | G | 383672 | 2.80 |

**>Ts7**
GCGGAUUUAmGCUCAGDDG

**Table S8.** 5´_biotin_tRNA_T1_SII_032919s07_44A45G. Sequencing of 5´ biotin labeled segment II from 21A to 57G by global hierarchical ranking algorithm.

| Fragment | Mass | RT | Base | Volume | PPM |
|---|---|---|---|---|---|
| 1 | 922.2241 | 25.229 | Tag+A | 745215 | 3.04 |
| 2 | 1267.2710 | 25.756 | G | 577150 | 2.60 |
| 3 | 1596.3229 | 28.405 | A | 472089 | 2.44 |
| 4 | 1941.3702 | 29.167 | G | 591742 | 2.06 |
| 5 | 2246.4125 | 30.221 | C | 930358 | 1.34 |
| 6 | 2619.4912 | 35.055 | 2mG | 276858 | 1.15 |
| 7 | 2924.5312 | 35.109 | C | 937840 | 1.47 |
| 8 | 3229.5745 | 35.989 | C | 1389357 | 0.71 |
| 9 | 3558.6244 | 37.535 | A | 944505 | 1.38 |
| 10 | 3903.6768 | 38.016 | G | 1334405 | 0.03 |
| 11 | 4232.7261 | 39.120 | A | 899666 | 0.73 |
| 12 | 4857.8097 | 40.778 | U+Cm | 2369525 | 0.37 |
| 13 | 5545.9261 | 42.941 | A+Gm | 1777156 | 0.18 |
| 14 | 5874.9889 | 43.512 | A | 1527490 | 1.60 |
| 15 | 6086.9945 | 43.461 | Y' | 2278504 | 1.05 |
| 16 | 6416.0477 | 44.268 | A | 1366254 | 1.11 |
| 17 | 6722.0827 | 44.327 | U | 1049995 | 2.48 |
| 18 | 7041.1313 | 44.591 | mC | 1297495 | 1.19 |
| 19 | 7347.1602 | 44.775 | U | 1560416 | 1.62 |
| 20 | 7692.2118 | 45.013 | G | 1319384 | 2.09 |
| 21 | 8037.2549 | 45.410 | G | 1009813 | 1.47 |
| 22 | 8366.3413 | 45.858 | A | 271843 | 5.46 |
| 23 | 8711.3823 | 45.865 | G | 1226283 | 4.51 |
| 24 | 9070.4677 | 45.822 | mG | 520562 | 6.79 |
| 25 | 9376.4389 | 45.871 | U | 416614 | 0.79 |
| 26 | 9681.5649 | 45.921 | C | 587268 | 9.51 |
| 27 | 10000.5521 | 46.069 | mC | 504658 | 2.24 |
| 28 | 10306.6258 | 46.099 | U | 925998 | 6.86 |
| 29 | 10651.5989 | 46.183 | G | 672326 | 0.34 |
| 30 | 10957.6318 | 46.200 | U | 320227 | 0.36 |
| 31 | 11302.6636 | 46.313 | G | 962623 | 1.04 |
| 32 | 11622.6493 | 46.492 | T | 325162 | 5.76 |
| 33 | 11928.6903 | 46.401 | U | 2182861 | 4.31 |
| 34 | 12233.7642 | 46.449 | C | 463444 | 1.54 |
| 35 | 12578.8603 | 46.548 | G | 2766678 | 2.38 |

>Ts8
AGAGC2mGCCAGACmUGmAAY´AUmCUGGAGmGUCmCUGUGTUCG

**Table S9.** 5´_biotin_tRNA_T1_SII_032919s07_44g45a. Sequencing of 5´ biotin labeled tRNA segment II from 21A to 57G by global hierarchical ranking algorithm.

| Fragment | Mass | RT | Base | Volume | PPM |
|---|---|---|---|---|---|
| 1 | 922.2241 | 25.229 | Tag+A | 745215 | 3.04 |
| 2 | 1267.2710 | 25.756 | G | 577150 | 2.60 |
| 3 | 1596.3229 | 28.405 | A | 472089 | 2.44 |
| 4 | 1941.3702 | 29.167 | G | 591742 | 2.06 |
| 5 | 2246.4125 | 30.221 | C | 930358 | 1.34 |
| 6 | 2619.4912 | 35.055 | 2mG | 276858 | 1.15 |
| 7 | 2924.5312 | 35.109 | C | 937840 | 1.47 |
| 8 | 3229.5745 | 35.989 | C | 1389357 | 0.71 |
| 9 | 3558.6244 | 37.535 | A | 944505 | 1.38 |
| 10 | 3903.6768 | 38.016 | G | 1334405 | 0.03 |
| 11 | 4232.7261 | 39.120 | A | 899666 | 0.73 |
| 12 | 4857.8097 | 40.778 | U+Cm | 2369525 | 0.37 |
| 13 | 5545.9261 | 42.941 | A+Gm | 1777156 | 0.18 |
| 14 | 5874.9889 | 43.512 | A | 1527490 | 1.58 |
| 15 | 6086.9945 | 43.461 | Y' | 2278504 | 1.03 |
| 16 | 6416.0477 | 44.268 | A | 1366254 | 1.09 |
| 17 | 6722.0827 | 44.327 | U | 1049995 | 2.48 |
| 18 | 7041.1313 | 44.591 | mC | 1297495 | 1.19 |
| 19 | 7347.1602 | 44.775 | U | 1560416 | 1.63 |
| 20 | 7692.2118 | 45.013 | G | 1319384 | 2.11 |
| 21 | 8037.2549 | 45.410 | G | 1009813 | 1.48 |
| 22 | 8382.2778 | 45.275 | G | 200964 | 1.50 |
| 23 | 8711.3823 | 45.865 | A | 1226283 | 4.53 |
| 24 | 9070.4677 | 45.822 | mG | 520562 | 6.80 |
| 25 | 9376.4389 | 45.871 | U | 416614 | 0.81 |
| 26 | 9681.5649 | 45.921 | C | 587268 | 9.53 |
| 27 | 10000.5521 | 46.069 | mC | 504658 | 2.26 |
| 28 | 10306.6258 | 46.099 | U | 925998 | 6.89 |
| 29 | 10651.5989 | 46.183 | G | 672326 | 0.31 |
| 30 | 10957.6318 | 46.200 | U | 320227 | 0.39 |
| 31 | 11302.6636 | 46.313 | G | 962623 | 1.00 |
| 32 | 11622.6493 | 46.492 | T | 325162 | 5.73 |
| 33 | 11928.6903 | 46.401 | U | 2182861 | 4.27 |
| 34 | 12233.7642 | 46.449 | C | 463444 | 1.50 |
| 35 | 12578.8603 | 46.548 | G | 2766678 | 2.42 |

>Ts9
AGAGC2mGCCAGACmUGmAAY´AUmCUGGGAmGUCmCUGUGTUCG

**Table S10.** 3´__tRNA_1009s06. Sequencing of acid degraded tRNA from 45G to 76A by global hierarchical ranking algorithm.

| Fragment | Mass | RT | Base | Volume | PPM |
|---|---|---|---|---|---|
| 1 | 877.1786 | 1.270 | A+C+C | 1022495 | 0.80 |
| 2 | 1206.2286 | 2.926 | A | 1172115 | 2.65 |
| 3 | 1511.2689 | 2.572 | C | 819385 | 2.78 |
| 4 | 1856.3153 | 3.218 | G | 1266301 | 2.80 |
| 5 | 2161.3551 | 3.798 | C | 1544446 | 3.10 |
| 6 | 2467.3789 | 4.806 | U | 2083726 | 3.36 |
| 7 | 2773.4034 | 5.685 | U | 1696734 | 3.32 |
| 8 | 3102.4553 | 7.075 | A | 5583907 | 3.16 |
| 9 | 3431.5054 | 7.910 | A | 2247902 | 3.56 |
| 10 | 3776.5516 | 7.745 | G | 5639286 | 3.55 |
| 11 | 4105.6016 | 8.447 | A | 2679354 | 3.87 |
| 12 | 4410.6408 | 8.523 | C | 4702025 | 4.08 |
| 13 | 4739.6917 | 9.123 | A | 2963739 | 4.14 |
| 14 | 5044.7319 | 9.175 | C | 2073512 | 4.10 |
| 15 | 5349.7949 | 9.288 | C | 1906782 | 0.19 |
| 16 | 5655.7967 | 9.545 | U | 914935 | 4.00 |
| 17 | 5998.8627 | 9.818 | mA | 2160204 | 4.12 |
| 18 | 6343.9049 | 9.900 | G | 2309111 | 4.71 |
| 19 | 6648.9464 | 9.893 | C | 3092250 | 4.47 |
| 20 | 6954.9754 | 9.838 | U | 1201050 | 3.75 |
| 21 | 7275.0127 | 10.396 | T | 2267279 | 4.10 |
| 22 | 7620.0765 | 10.498 | G | 1762814 | 1.76 |
| 23 | 7926.1455 | 10.423 | U | 1562423 | 3.81 |
| 24 | 8271.1067 | 10.603 | G | 1920966 | 6.77 |
| 25 | 8577.2011 | 10.660 | U | 1709835 | 1.52 |
| 26 | 8896.1598 | 11.550 | mC | 875226 | 9.58 |
| 27 | 9201.2581 | 11.313 | C | 769527 | 3.06 |
| 28 | 9507.2765 | 11.082 | U | 572956 | 3.70 |
| 29 | 9866.3028 | 11.030 | mG | 412887 | 7.30 |
| 30 | 10211.3522 | 11.073 | G | 709961 | 6.86 |

>Ts10
GmGUCmCUGUGTUCGmAUCCACAGAAUUCGCACCA

**Table S11.** 5´_pG_tRNA_100918s06. Sequencing of 5´pG tRNA from 1G to 31A by global hierarchical ranking algorithm.

| Fragment | Mass | RT | Base | Volume | PPM |
|---|---|---|---|---|---|
| 1 | 443.0274 | 0.931 | pG | 233231 | 7.00 |
| 2 | 748.0684 | 1.039 | C | 883929 | 3.74 |
| 3 | 1093.1105 | 1.800 | G | 2062278 | 2.29 |
| 4 | 1438.1575 | 3.239 | G | 3687690 | 2.02 |
| 5 | 1767.2087 | 4.484 | A | 4522172 | 2.38 |
| 6 | 2073.2354 | 5.369 | U | 8131266 | 1.35 |
| 7 | 2379.2590 | 6.043 | U | 8862830 | 1.89 |
| 8 | 2685.2836 | 6.593 | U | 9612100 | 1.94 |
| 9 | 3014.3343 | 7.355 | A | 6218090 | 2.32 |
| 10 | 3373.3964 | 8.120 | mG | 2974994 | 2.37 |
| 11 | 3678.4380 | 8.403 | C | 3957178 | 2.09 |
| 12 | 3984.4601 | 8.709 | U | 6419872 | 2.74 |
| 13 | 4289.5007 | 8.942 | C | 8348561 | 2.70 |
| 14 | 4618.5517 | 9.346 | A | 3797284 | 2.84 |
| 15 | 4963.6043 | 9.522 | G | 217686 | 1.59 |
| 16 | 5271.6374 | 9.631 | D | 3108073 | 3.00 |
| 17 | 5579.6773 | 9.748 | D | 3781679 | 3.03 |
| 18 | 5924.7327 | 9.944 | G | 689750 | 1.50 |
| 19 | 6269.7714 | 10.091 | G | 2753572 | 2.81 |
| 20 | 6614.8124 | 10.232 | G | 1506355 | 3.63 |
| 21 | 6943.8650 | 10.468 | A | 1708708 | 3.44 |
| 22 | 7288.9012 | 10.601 | G | 779104 | 4.82 |
| 23 | 7617.9417 | 10.826 | A | 852001 | 6.18 |
| 24 | 7963.0075 | 10.910 | G | 2445671 | 3.60 |
| 25 | 8268.0027 | 11.143 | C | 1087860 | 9.05 |
| 26 | 8641.1310 | 11.694 | 2mG | 207499 | 2.92 |
| 27 | 8946.1664 | 11.727 | C | 1364582 | 3.48 |
| 28 | 9251.2074 | 11.743 | C | 1059830 | 3.39 |
| 29 | 9580.2455 | 11.864 | A | 1450228 | 4.78 |
| 30 | 9925.3349 | 11.871 | G | 2494820 | 0.38 |
| 31 | 10254.2927 | 11.993 | A | 155606 | 9.61 |

>Ts11
GCGGAUUUAmGCUCAGDDGGGAGAGC2mGCCAGA

**Table S12.** Yield of CMC conversion occurring at pseudouridine measured by LC-MS.

| Conversion state | Fragment | Calc mass | Exp mass | m/z | EIC | QS | ppm |
|---|---|---|---|---|---|---|---|
| Non-converted | 21A to 44A | 7791.1320 | 7791.1787 | 778.1111 | 1129053 | 80 | -5.99 |
| CMC-converted | 21A to 44A | 8042.3318 | 8042.3492 | 803.2263 | 4123573 | 80 | -2.16 |
| Non-converted | 57G to 47U | 3526.4344 | 3526.4333 | 586.7314 | 1176461 | 100 | 0.31 |
| CMC-converted | 57G to 47A | 3777.6342 | 3777.6332 | 628.5979 | 3779411 | 100 | 0.26 |

**Table S13.** 5´_tRNA_T1_nonCMC_SII_042519s04_44A45G. Sequencing of 5´ non-CMC converted tRNA segment II from 21A to 45G by global hierarchical ranking algorithm.

| Fragment | Mass | RT | Base | Volume | PPM |
|---|---|---|---|---|---|
| 1 | 692.1076 | 1.032 | A+G | 121835 | 4.19 |
| 2 | 1021.1576 | 1.264 | A | 548483 | 5.29 |
| 3 | 1366.2072 | 4.020 | G | 2219430 | 2.34 |
| 4 | 1671.2480 | 7.304 | C | 3142702 | 2.21 |
| 5 | 2044.3269 | 16.800 | 2mG | 1700693 | 1.71 |
| 6 | 2349.3689 | 18.430 | C | 2431764 | 1.19 |
| 7 | 2654.4105 | 20.727 | C | 6691067 | 0.94 |
| 8 | 2983.4639 | 23.756 | A | 9276684 | 0.54 |
| 9 | 3328.5120 | 25.192 | G | 10673175 | 0.27 |
| 10 | 3657.5668 | 27.417 | A | 5126136 | 0.38 |
| 11 | 4282.6486 | 30.874 | U+Cm | 15880661 | 0.21 |
| 12 | 4970.7665 | 34.609 | A+Gm | 10873309 | 0.64 |
| 13 | 5299.8210 | 35.684 | A | 12807606 | 0.98 |
| 14 | 5511.8306 | 35.900 | Y' | 13088146 | 1.12 |
| 15 | 5840.8850 | 37.167 | A | 3623732 | 1.39 |
| 16 | 6146.9096 | 37.460 | U | 1897334 | 1.20 |
| 17 | 6465.9704 | 38.006 | mC | 2463925 | 1.75 |
| 18 | 6771.9928 | 38.393 | U | 3706693 | 1.24 |
| 19 | 7117.0453 | 38.873 | G | 3506106 | 1.90 |
| 20 | 7462.0964 | 39.527 | G | 2455794 | 2.30 |
| 21 | 7791.1787 | 40.196 | A | 1226259 | 6.03 |
| 22 | 8136.1916 | 40.385 | G | 1925167 | 1.54 |

>Ts13
AGAGC2mGCCAGACmUGmAAY´AUmCUGGAG

**Table S14**. 5´_tRNA_T1_nonCMC_SII_042519s04_44g45a. Sequencing of 5´ non-CMC converted tRNA segment II from 21A to 45A by global hierarchical ranking algorithm.

| Fragment | Mass | RT | Base | Volume | PPM |
|---|---|---|---|---|---|
| 1 | 692.1076 | 1.032 | A+G | 121835 | 4.19 |
| 2 | 1021.1576 | 1.264 | A | 548483 | 5.29 |
| 3 | 1366.2072 | 4.020 | G | 2219430 | 2.34 |
| 4 | 1671.2480 | 7.304 | C | 3142702 | 2.21 |
| 5 | 2044.3269 | 16.800 | 2mG | 1700693 | 1.71 |
| 6 | 2349.3689 | 18.430 | C | 2431764 | 1.19 |
| 7 | 2654.4105 | 20.727 | C | 6691067 | 0.94 |
| 8 | 2983.4639 | 23.756 | A | 9276684 | 0.54 |
| 9 | 3328.5120 | 25.192 | G | 10673175 | 0.27 |
| 10 | 3657.5668 | 27.417 | A | 5126136 | 0.38 |
| 11 | 4282.6486 | 30.874 | U+Cm | 15880661 | 0.21 |
| 12 | 4970.7665 | 34.609 | A+Gm | 10873309 | 0.64 |
| 13 | 5299.8210 | 35.684 | A | 12807606 | 0.98 |
| 14 | 5511.8306 | 35.900 | Y' | 13088146 | 1.12 |
| 15 | 5840.8850 | 37.167 | A | 3623732 | 1.39 |
| 16 | 6146.9096 | 37.460 | U | 1897334 | 1.20 |
| 17 | 6465.9704 | 38.006 | mC | 2463925 | 1.75 |
| 18 | 6771.9928 | 38.393 | U | 3706693 | 1.24 |
| 19 | 7117.0453 | 38.873 | G | 3506106 | 1.90 |
| 20 | 7462.0964 | 39.527 | G | 2455794 | 2.30 |
| 21 | 7807.1385 | 39.523 | G | 835117 | 1.52 |
| 22 | 8136.1916 | 40.385 | A | 1925167 | 1.54 |

>Ts14
AGAGC2mGCCAGACmUGmAAY´AUmCUGGGA

**Table S15.** 5´_tRNA_T1_CMC_SII_042519s04. Sequencing of 5´ CMC converted tRNA segment II from 39ψ to 44A by global hierarchical ranking algorithm.

| Fragment | Mass | RT | Base | Volume | PPM |
|---|---|---|---|---|---|
| 1 | 6398.1211 | 44.707 | Mod-Psi | 1295323 | 2.97 |
| 2 | 6717.1789 | 45.223 | mC | 2506731 | 2.96 |
| 3 | 7023.1878 | 45.283 | U | 3037253 | 0.50 |
| 4 | 7368.2361 | 45.446 | G | 8115206 | 0.60 |
| 5 | 7713.3006 | 45.574 | G | 4221938 | 2.79 |
| 6 | 8042.3492 | 46.255 | A | 3190026 | 2.19 |

>Ts15
ψmCUGGA

**Table S16.** 3´_tRNA_T1_nonCMC_SII_042519s04. Sequencing of 3´ non-CMC converted tRNA segment II from 57G to 47U by global hierarchical ranking algorithm.

| Fragment | Mass | RT | Base | Volume | PPM |
|---|---|---|---|---|---|
| 1 | 668.0943 | 0.968 | G+C | 79549 | 7.33 |
| 2 | 974.1302 | 0.915 | U | 826458 | 5.85 |
| 3 | 1294.1594 | 2.732 | T | 403523 | 4.71 |
| 4 | 1639.2089 | 6.500 | G | 789168 | 2.44 |
| 5 | 1945.2357 | 6.129 | U | 190380 | 1.29 |
| 6 | 2290.2818 | 10.466 | G | 1584520 | 1.66 |
| 7 | 2596.3069 | 12.965 | U | 1100858 | 1.54 |
| 8 | 2915.3646 | 17.907 | mC | 1557574 | 1.10 |
| 9 | 3220.4052 | 18.523 | C | 773618 | 1.21 |
| 10 | 3526.4333 | 20.318 | U | 2252901 | 0.31 |

>Ts16
UCmCUGUGTUCG

**Table S17.** 3´_tRNA_T1_CMC_SII_042519s04. Sequencing of 3´ CMC converted tRNA segment II from 57G to 47U by global hierarchical ranking algorithm.

| Fragment | Mass | RT | Base | Volume | PPM |
|---|---|---|---|---|---|
| 1 | 1225.3215 | 14.484 | Mod-Psi | 882395 | 2.29 |
| 2 | 1545.3611 | 19.764 | T | 78086 | 2.72 |
| 3 | 1890.4097 | 27.200 | G | 1324986 | 1.59 |
| 4 | 2196.4340 | 25.561 | U | 33874 | 1.82 |
| 5 | 2541.4824 | 27.899 | G | 3029272 | 1.18 |
| 6 | 2847.5087 | 28.729 | U | 2275337 | 0.70 |
| 7 | 3166.5661 | 32.358 | mC | 2499558 | 0.47 |
| 8 | 3471.6055 | 32.073 | C | 2485944 | 0.98 |
| 9 | 3777.6332 | 32.777 | U | 4553148 | 0.26 |

>Ts17
UCmCUGUGTψ

**Table S18.** Detection of Y´ in the presence of tRNA before (in full-length tRNA) and after (as isolated base) acid degradation.

| In a form of segment II | Calc mass | Exp mass | m/z | EIC | Percent | QS | ppm |
|---|---|---|---|---|---|---|---|
| Y before acid degradation | 12361.805 | 12361.841 | 823.1141 | 2324857 | 90% | 80 | -2.9 |
| Y´ before acid degradation | 12003.666 | 12003.762 | 922.359 | 230727 | 10% | 48 | -7.9 |
| Y´ after acid degradation | 376.1495 | 376.1479 | 375.1409 | 49059213 | 100% | 100 | 4.3 |

**Table S19.** 5´_OH_tRNA_T1_SII_111418s05_44A45G. LC-MS analysis of segment II from 34Gm to 55ψ (mass ladder components from 3´ to 5´). The sequence was manually verified.

| Theoretical | | | | Extracted data file after LC/MS analysis | | | | Error |
|---|---|---|---|---|---|---|---|---|
| Fragments | Theoretical mass | Base mass | Base | MFE mass | $t_R$ | Volume | Quality Score | ppm |
| 21 | 7739.0291 | 688.1156 | A+Gm | 7739.0198 | 28.919 | 572629 | 80 | 1.20 |
| 20 | 7050.9135 | 329.0525 | A | 7050.9277 | 26.539 | 413840 | 60 | -2.01 |
| 19 | 6721.8610 | 212.0086 | Y´ | 6721.8635 | 24.741 | 381223 | 72.8 | -0.37 |
| 18 | 6509.8524 | 329.0525 | A | 6509.8604 | 25.336 | 1019699 | 80 | -1.23 |
| 17 | 6180.7999 | 306.0253 | ψ | 6180.8037 | 23.079 | 707995 | 77.8 | -0.61 |
| 16 | 5874.7746 | 319.0570 | $m^5C$ | 5874.7783 | 23.641 | 2167527 | 100 | -0.63 |
| 15 | 5555.7176 | 306.0253 | U | 5555.7209 | 21.539 | 1146864 | 98.5 | -0.59 |
| 14 | 5249.6923 | 345.0474 | G | 5249.6958 | 20.605 | 1609784 | 100 | -0.67 |
| 13 | 4904.6449 | 345.0475 | G | 4904.6446 | 19.764 | 1791176 | 100 | 0.06 |
| 12 | 4559.5974 | 329.0525 | A | 4559.5918 | 19.341 | 974223 | 80 | 1.23 |
| 11 | 4230.5449 | 345.0474 | G | 4230.5449 | 16.828 | 1254040 | 99.7 | 0.00 |
| 10 | 3885.4975 | 359.0631 | $m^7G$ | 3885.4957 | 15.319 | 1940572 | 95.7 | 0.46 |
| 9 | 3526.4344 | 306.0253 | U | 3526.4327 | 13.475 | 1011995 | 100 | 0.48 |
| 8 | 3220.4091 | 305.0413 | C | 3220.4066 | 11.393 | 2082145 | 100 | 0.78 |
| 7 | 2915.3678 | 319.0569 | $m^5C$ | 2915.3648 | 10.586 | 3108932 | 100 | 1.03 |
| 6 | 2596.3109 | 306.0253 | U | 2596.3066 | 6.488 | 523377 | 42.8 | 1.66 |
| 5 | 2290.2856 | 345.0475 | G | 2290.2828 | 3.961 | 2464626 | 94.7 | 1.22 |
| 4 | 1945.2381 | 306.0253 | U | 1945.2379 | 1.074 | 637786 | 83.4 | 0.10 |
| 3 | 1639.2128 | 345.0474 | G | 1639.2106 | 1.034 | 2301078 | 100 | 1.34 |
| 2 | 1294.1654 | 320.0409 | T | 1294.1737 | 8.127 | 78112 | 67.5 | -6.41 |
| 1 | 974.1245 | 306.0253 | ψ | 974.1240 | 0.936 | 143886 | 79.1 | 0.51 |

>Ts19
GmAAY´AUmCUGGAGmGUCmCUGUGTU

**Table S20.** 5´_OH_tRNA_T1_SII_111418s05_44g45a. LC-MS analysis of segment II from 34Gm to 55ψ (mass ladder components from 3´ to 5´). The sequence was manually verified.

| **Theoretical** | | | | **Extracted data file after LC/MS analysis** | | | | **Error** |
|---|---|---|---|---|---|---|---|---|
| **Fragments** | **Theoretical mass** | **Base mass** | **Base** | **MFE mass** | **$t_R$** | **Volume** | **Quality Score** | **ppm** |
| 21 | 7739.0291 | 688.1156 | A+Gm | 7739.0198 | 28.919 | 572629 | 80 | 1.20 |
| 20 | 7050.9135 | 329.0525 | A | 7050.9277 | 26.539 | 413840 | 60 | -2.01 |
| 19 | 6721.8610 | 212.0086 | Y´ | 6721.8635 | 24.741 | 381223 | 72.8 | -0.37 |
| 18 | 6509.8524 | 329.0525 | A | 6509.8604 | 25.336 | 1019699 | 80 | -1.23 |
| 17 | 6180.7999 | 306.0253 | ψ | 6180.8037 | 23.079 | 707995 | 77.8 | -0.61 |
| 16 | 5874.7746 | 319.0570 | $m^5C$ | 5874.7783 | 23.641 | 2167527 | 100 | -0.63 |
| 15 | 5555.7176 | 306.0253 | U | 5555.7209 | 21.539 | 1146864 | 98.5 | -0.59 |
| 14 | 5249.6923 | 345.0474 | G | 5249.6958 | 20.605 | 1609784 | 100 | -0.67 |
| 13 | 4904.6449 | 345.0475 | G | 4904.6446 | 19.764 | 1791176 | 100 | 0.06 |
| 12 | 4559.5974 | 345.0474 | G | 4559.5918 | 19.341 | 974223 | 80 | 100 |
| 11 | 4214.5500 | 329.0525 | A | 4214.5624 | 18.424 | 273170 | 79.6 | 100 |
| 10 | 3885.4975 | 359.0631 | $m^7G$ | 3885.4957 | 15.319 | 1940572 | 95.7 | 0.46 |
| 9 | 3526.4344 | 306.0253 | U | 3526.4327 | 13.475 | 1011995 | 100 | 0.48 |
| 8 | 3220.4091 | 305.0413 | C | 3220.4066 | 11.393 | 2082145 | 100 | 0.78 |
| 7 | 2915.3678 | 319.0569 | $m^5C$ | 2915.3648 | 10.586 | 3108932 | 100 | 1.03 |
| 6 | 2596.3109 | 306.0253 | U | 2596.3066 | 6.488 | 523377 | 42.8 | 1.66 |
| 5 | 2290.2856 | 345.0475 | G | 2290.2828 | 3.961 | 2464626 | 94.7 | 1.22 |
| 4 | 1945.2381 | 306.0253 | U | 1945.2379 | 1.074 | 637786 | 83.4 | 0.10 |
| 3 | 1639.2128 | 345.0474 | G | 1639.2106 | 1.034 | 2301078 | 100 | 1.34 |
| 2 | 1294.1654 | 320.0409 | T | 1294.1737 | 8.127 | 78112 | 67.5 | -6.41 |
| 1 | 974.1245 | 306.0253 | ψ | 974.1240 | 0.936 | 143886 | 79.1 | 0.51 |

>Ts20
GmAAY´AUmCUGGGAmGUCmCUGUGTU

**Table S21.** 5´_biotin_tRNA_T1_SII_032919s07_44A45G. LC-MS analysis of segment II from 30G to 55ψ (mass ladder components from 3´ to 5´). The sequence was manually verified.

| Theoretical | | | | Extracted data file after LC/MS analysis | | | | Error |
|---|---|---|---|---|---|---|---|---|
| **Fragments** | **Theoretical mass** | **Base mass** | **Base** | **MFE mass** | **$t_R$** | **Volume** | **Quality Score** | **ppm** |
| 24 | 9038.2113 | 345.0474 | G | 9038.133 | 37.926 | 394860 | 60.8 | 8.66 |
| 23 | 8693.1639 | 329.0525 | A | 8693.1871 | 38.113 | 174673 | 41.4 | -2.67 |
| 22 | 8364.1114 | 625.0823 | U+Cm | 8364.1502 | 37.005 | 133633 | 41.9 | -4.64 |
| 21 | 7739.0291 | 688.1156 | A+Gm | 7739.0557 | 35.391 | 650792 | 77.4 | -3.44 |
| 20 | 7050.9135 | 329.0525 | A | 7050.9339 | 32.627 | 590137 | 78.5 | -2.89 |
| 19 | 6721.8610 | 212.0086 | Y´ | 6721.8845 | 30.813 | 764391 | 80 | -3.50 |
| 18 | 6509.8524 | 329.0525 | A | 6509.864 | 31.762 | 1166876 | 80 | -1.78 |
| 17 | 6180.7999 | 306.0253 | ψ | 6180.7968 | 29.159 | 148437 | 65.9 | 0.50 |
| 16 | 5874.7746 | 319.0570 | $m^5C$ | 5874.7784 | 30.31 | 1368105 | 79.9 | -0.65 |
| 15 | 5555.7176 | 306.0253 | U | 5555.7219 | 27.737 | 1148576 | 80 | -0.77 |
| 14 | 5249.6923 | 345.0474 | G | 5249.7098 | 26.957 | 1297236 | 80 | -3.33 |
| 13 | 4904.6449 | 345.0475 | G | 4904.6497 | 26.195 | 1021939 | 90 | -0.98 |
| 12 | 4559.5974 | 329.0525 | A | 4559.5974 | 25.942 | 1209559 | 99 | 0.00 |
| 11 | 4230.5449 | 345.0474 | G | 4230.5461 | 23.338 | 927818 | 92.3 | -0.28 |
| 10 | 3885.4975 | 359.0631 | $m^7G$ | 3885.4975 | 21.811 | 1357508 | 90.5 | 0.00 |
| 9 | 3526.4344 | 306.0253 | U | 3526.4332 | 20.034 | 1078413 | 98.3 | 0.34 |
| 8 | 3220.4091 | 305.0413 | C | 3220.4063 | 18.209 | 1434999 | 100 | 0.87 |
| 7 | 2915.3678 | 319.0569 | $m^5C$ | 2915.366 | 17.589 | 2388681 | 100 | 0.62 |
| 6 | 2596.3109 | 306.0253 | U | 2596.308 | 12.655 | 1592241 | 100 | 1.12 |
| 5 | 2290.2856 | 345.0475 | G | 2290.2828 | 10.189 | 2053112 | 100 | 1.22 |
| 4 | 1945.2381 | 306.0253 | U | 1945.2371 | 6.47 | 1359480 | 77.8 | 0.51 |
| 3 | 1639.2128 | 345.0474 | G | 1639.21 | 4.723 | 1598482 | 100 | 1.71 |
| 2 | 1294.1654 | 320.0409 | T | 1294.1615 | 2.282 | 620026 | 100 | 3.01 |
| 1 | 974.1245 | 306.0253 | ψ | 974.1225 | 0.875 | 221837 | 90.6 | 2.05 |

>Ts21
GACmUGmAAY´AUmCUGGAGmGUCmCUGUGTU

**Table S22.** 5´_biotin_tRNA_T1_SII_032919s07_44g45a. LC-MS analysis of segment II from 30G to 55ψ (mass ladder components from 3´ to 5´). The sequence was manually verified.

| **Theoretical** | | | | **Extracted data file after LC/MS analysis** | | | | **Error** |
|---|---|---|---|---|---|---|---|---|
| **Fragments** | **Theoretical mass** | **Base mass** | **Base** | **MFE mass** | **$t_R$** | **Volume** | **Quality Score** | **ppm** |
| 24 | 9038.2113 | 345.0474 | G | 9038.133 | 37.926 | 394860 | 60.8 | 8.66 |
| 23 | 8693.1639 | 329.0525 | A | 8693.1871 | 38.113 | 174673 | 41.4 | -2.67 |
| 22 | 8364.1114 | 625.0823 | U+Cm | 8364.1502 | 37.005 | 133633 | 41.9 | -4.64 |
| 21 | 7739.0291 | 688.1156 | A+Gm | 7739.0557 | 35.391 | 650792 | 77.4 | -3.44 |
| 20 | 7050.9135 | 329.0525 | A | 7050.9339 | 32.627 | 590137 | 78.5 | -2.89 |
| 19 | 6721.8610 | 212.0086 | Y´ | 6721.8845 | 30.813 | 764391 | 80 | -3.50 |
| 18 | 6509.8524 | 329.0525 | A | 6509.864 | 31.762 | 1166876 | 80 | -1.78 |
| 17 | 6180.7999 | 306.0253 | ψ | 6180.7968 | 29.159 | 148437 | 65.9 | 0.50 |
| 16 | 5874.7746 | 319.0570 | $m^5C$ | 5874.7784 | 30.31 | 1368105 | 79.9 | -0.65 |
| 15 | 5555.7176 | 306.0253 | U | 5555.7219 | 27.737 | 1148576 | 80 | -0.77 |
| 14 | 5249.6923 | 345.0474 | G | 5249.7098 | 26.957 | 1297236 | 80 | -3.33 |
| 13 | 4904.6449 | 345.0475 | G | 4904.6497 | 26.195 | 1021939 | 90 | -0.98 |
| 12 | 4559.5974 | 345.0474 | G | 4559.5974 | 25.942 | 1209559 | 99 | 0.00 |
| 11 | 4214.5500 | 329.0525 | A | 4214.5534 | 24.918 | 299777 | 60 | -0.81 |
| 10 | 3885.4975 | 359.0631 | $m^7G$ | 3885.4975 | 21.811 | 1357508 | 90.5 | 0.00 |
| 9 | 3526.4344 | 306.0253 | U | 3526.4332 | 20.034 | 1078413 | 98.3 | 0.34 |
| 8 | 3220.4091 | 305.0413 | C | 3220.4063 | 18.209 | 1434999 | 100 | 0.87 |
| 7 | 2915.3678 | 319.0569 | $m^5C$ | 2915.366 | 17.589 | 2388681 | 100 | 0.62 |
| 6 | 2596.3109 | 306.0253 | U | 2596.308 | 12.655 | 1592241 | 100 | 1.12 |
| 5 | 2290.2856 | 345.0475 | G | 2290.2828 | 10.189 | 2053112 | 100 | 1.22 |
| 4 | 1945.2381 | 306.0253 | U | 1945.2371 | 6.47 | 1359480 | 77.8 | 0.51 |
| 3 | 1639.2128 | 345.0474 | G | 1639.21 | 4.723 | 1598482 | 100 | 1.71 |
| 2 | 1294.1654 | 320.0409 | T | 1294.1615 | 2.282 | 620026 | 100 | 3.01 |
| 1 | 974.1245 | 306.0253 | ψ | 974.1225 | 0.875 | 221837 | 90.6 | 2.05 |

>Ts22
GACmUGmAAY´AUmCUGGGAmGUCmCUGUGTU

**Table S23.** Detection of wild type (44A45G) and transition form (44g45a), respectively, in three datasets by global hierarchical ranking algorithm (refer to output files Table S4, 5, 8, 9, 13 and 14).

| Dataset | Wild type (I) | | | | Transition form (II) | | | | I % | Mean± SEM | II % | Mean± SEM |
|---|---|---|---|---|---|---|---|---|---|---|---|---|
| | m/z | EIC (44A) | m/z | EIC (45G) | m/z | EIC (44G) | m/z | EIC (45A) | | | | |
| Labeled segment II | 836.1243 | 2308326 | 870.4306 | 1994979 | 837.6269 | 1932380 | 870.4306 | 1994979 | 54 | 50.4±3.2% | 46 | 49.6±3.2% |
| Unlabeled segment II | 778.4074 | 2077840 | 812.9122 | 1608093 | 780.0080 | 2630985 | 812.9122 | 1608093 | 44 | | 56 | |
| Non-CMC-converted segment II | 778.4077 | 1385023 | 813.0133 | 1770337 | 779.7066 | 1245805 | 813.0133 | 1770337 | 53 | | 47 | |

*Form I % = EIC(44A) / EIC(44A) + EIC(44G); Form II % = EIC(44G) / EIC(44A) + EIC(44G)
